# Structural basis of the activation of the CC chemokine receptor 5 by a chemokine agonist

**DOI:** 10.1101/2020.11.27.401117

**Authors:** Polina Isaikina, Ching-Ju Tsai, Nikolaus Dietz, Filip Pamula, Anne Grahl, Kenneth N. Goldie, Ramon Guixà-González, Gebhard F.X. Schertler, Oliver Hartley, Henning Stahlberg, Timm Maier, Xavier Deupi, Stephan Grzesiek

**Author notes:** Address correspondence to: Stephan Grzesiek, Focal Area Structural Biology and Biophysics, Biozentrum, University of Basel, CH-4056 Basel, Switzerland, Xavier Deupi, Timm Maier, Oliver Hartley, Gebhard F.X. Schertler.

## Abstract

The human CC chemokine receptor 5 (CCR5) is a G protein-coupled receptor (GPCR) that plays a major role in inflammation and is involved in the pathology of cancer, HIV, and COVID-19. Despite its significance as a drug target, the activation mechanism of CCR5, i.e. how chemokine agonists transduce the activation signal through the receptor, is yet unknown. Here, we report the cryo-EM structure of wild-type CCR5 in an active conformation bound to the chemokine super-agonist [6P4]CCL5 and the heterotrimeric G_i_ protein. The structure provides the rationale for the sequence-activity relation of agonist and antagonist chemokines. The N-terminus of agonist chemokines pushes onto an aromatic connector that transmits activation to the canonical GPCR microswitch network. This activation mechanism differs significantly from other CC chemokine receptors that bind shorter chemokines in a shallow binding mode and have unique sequence signatures and a specialized activation mechanism.

**One-sentence summary:** The structure of CCR5 in complex with the chemokine agonist [6P4]CCL5 and the heterotrimeric Gi protein reveals its activation mechanism

---

The human CC chemokine receptor 5 (CCR5) is a G protein-coupled receptor (GPCR) that plays a major role in inflammation by recruiting and activating leukocytes(*1*). CCR5 is also the principal HIV coreceptor(*2*), is involved in the pathology of both cancer(*3*) and neuroinflammation(*4*), and has been implicated in the inflammatory complications of COVID-19(*5, 6*). Soon after the discovery of CCR5, it became evident that its natural chemokine ligands inhibit HIV entry(*7*), with CCL5 (RANTES) being most efficient, both by blocking the binding site for the viral glycoprotein gp120 and by promoting CCR5 endocytosis(*8*). Modifications of the N-terminal region of CCR5 preceding C10 yielded HIV entry inhibitors with significantly higher potency(*9–11*). These analogs belong to a group of over 100 engineered CCL5 N-terminal variants that show striking differences in their anti-HIV, endocytotic, affinity, and signaling properties ranging e.g. from super-agonist to strong antagonist behavior(*10, 11*). The molecular explanation for these N-terminal structure-related differences is currently unclear.

Whereas a good structural understanding has been reached of the activation mechanisms of class A GPCRs by small-molecule ligands(*12*), the activation mechanism of the chemokine receptor subclass is not yet well understood. Inactive structures of a number of chemokine receptors have been solved including complexes of CCR5 with the engineered chemokine antagonist [5P7]CCL5(*13*), the viral gp120•human CD4 complex(*14*), the HIV inhibitor maraviroc(*15*), and other small-molecule antagonists(*16*). In contrast, currently only three active-state chemokine receptor complex structures are available: CX3CL1•US28•Nb7(*17*), CCL20•CCR6•G_o_(*18*), and CXCL8•CXCR2•G_i_(*19*). The viral receptor US28 is constitutively active and hence does not reveal chemokine-induced activation, and the chemokines CCL20 and CXCL8 adopt a shallow binding mode in which activation apparently involves transmission of forces directly from the extracellular domain of the receptor. On the other hand, receptors such as CCR5 have ligands with longer N-termini which likely insert more deeply into the orthosteric pocket of the receptor (chemokine recognition site 2; CRS2). At least in the case of CCR5, where agonist and antagonist N-terminal CCL5 variants of the same length exist, crucial signaling interactions must rather be located within CRS2.

With the aim of elucidating the apparently different activation mechanisms of CC chemokine receptors and to provide a general structural explanation for the variable pharmacology of CCL5 N-terminal variants, we solved the structure of wild-type human CCR5 in complex with the super-agonist [6P4]CCL5 and the G_i_ heterotrimer.

## Overall structure of the [6P4]CCL5•CCR5•G_i_ complex

A stable complex of [6P4]CCL5•CCR5•G_i_ was obtained by incubating detergent-solubilized human wild-type full-length CCR5 with the G_i1_ heterotrimer and [6P4]CCL5 (Supplementary Fig. S1). The complex was treated with apyrase to hydrolyze GDP and was further stabilized by addition of the Fab fragment Fab16(*20, 21*), which recognizes an interface between the Gα and Gβγ subunits of the G_i_ heterotrimer (Supplementary Fig. S2). Single-particle cryo-electron microscopy (cryo-EM) analysis with extensive particle classification yielded a threedimensional density map with a nominal global resolution of 3.1 Å (Fig. 1A, Supplementary Fig. S3, and Supplementary Table S1). The map is well resolved for most of parts of CCR5, the [6P4]CCL5 N-terminus, the G_i_ heterotrimer, and Fab16 (Supplementary Fig. S4). The density of the globular core of [6P4]CCL5 and the adjacent CCR5 N-terminus and extracellular parts of the receptor have less-defined density, indicating relative flexibility in these parts of the structure. Indeed, a 3D variability analysis of the cryo-EM data (Supplementary Video S1) and molecular dynamics simulations of the atomic model (Supplementary Fig. S5A) reveal a certain degree of mobility of the [6P4]CCL5 core as well as of the receptor N-terminus, extracellular loops, TM5, 6 and 7. Apart from a small ~5° difference in the orientation, the position of the [6P4]CCL5 core is very similar to that of [5P7]CCL5 in the inactive [5P7]CCL5•CCR5 complex (Fig. 1C, 2A). Nevertheless, this minor change in orientation also leads to small (1–2 Å), but noticeable movements at the extracellular helix ends (Fig. 2B).

**Figure 1.**
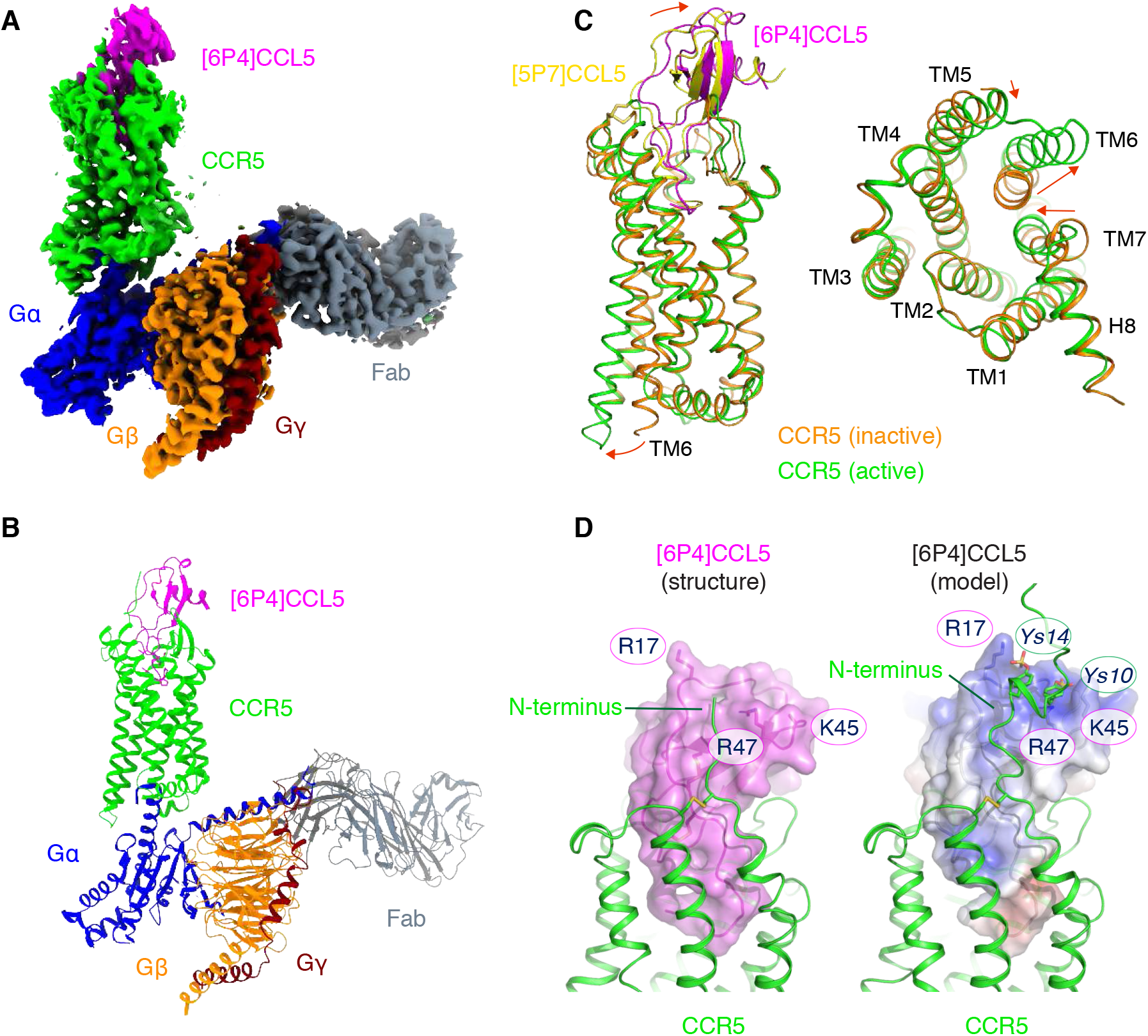
Cryo-EM structure of the [6P4]CCL5•CCR5•G_i_•Fab16 complex. (A) Cryo-EM map of the [6P4]CCL5•CCR5•G_i_•Fab16 complex colored by subunits ([6P4]CCL5 – magenta, CCR5 – green, Gα_i_ – blue, Gβ – orange, Gγ – maroon, Fab16 – grey). (B) Atomic model of the [6P4]CCL5•CCR5•G_i_•Fab16 complex in the same view and color scheme as shown in (A). (C) Side and cytoplasmic views of the structural overlay of active CCR5 (green) in complex [6P4]CCL5 (magenta) and inactive CCR5 (orange, PDB ID: 5UIW) in complex with [5P7]CCL5 (yellow). Significant structural changes between two conformations are indicated by red arrows. The C101^3.25^-C178^ECL2^, C20^N-term^-C269^7.25^ disulphide bridges conserved chemokine receptors are shown in dark yellow. (D) Interactions between the [6P4]CCL5 core and the CCR5 N-terminus at the CRS1 site (left, cryo-EM structure; right: combined cryo-EM/NMR model). In the model, sulfo-tyrosines ^s^Y10 and ^s^Y14 are depicted as sticks and the [6P4]CCL5 surface is colored according to its electrostatic potential (negative: red; positive: blue).

**Figure 2.**
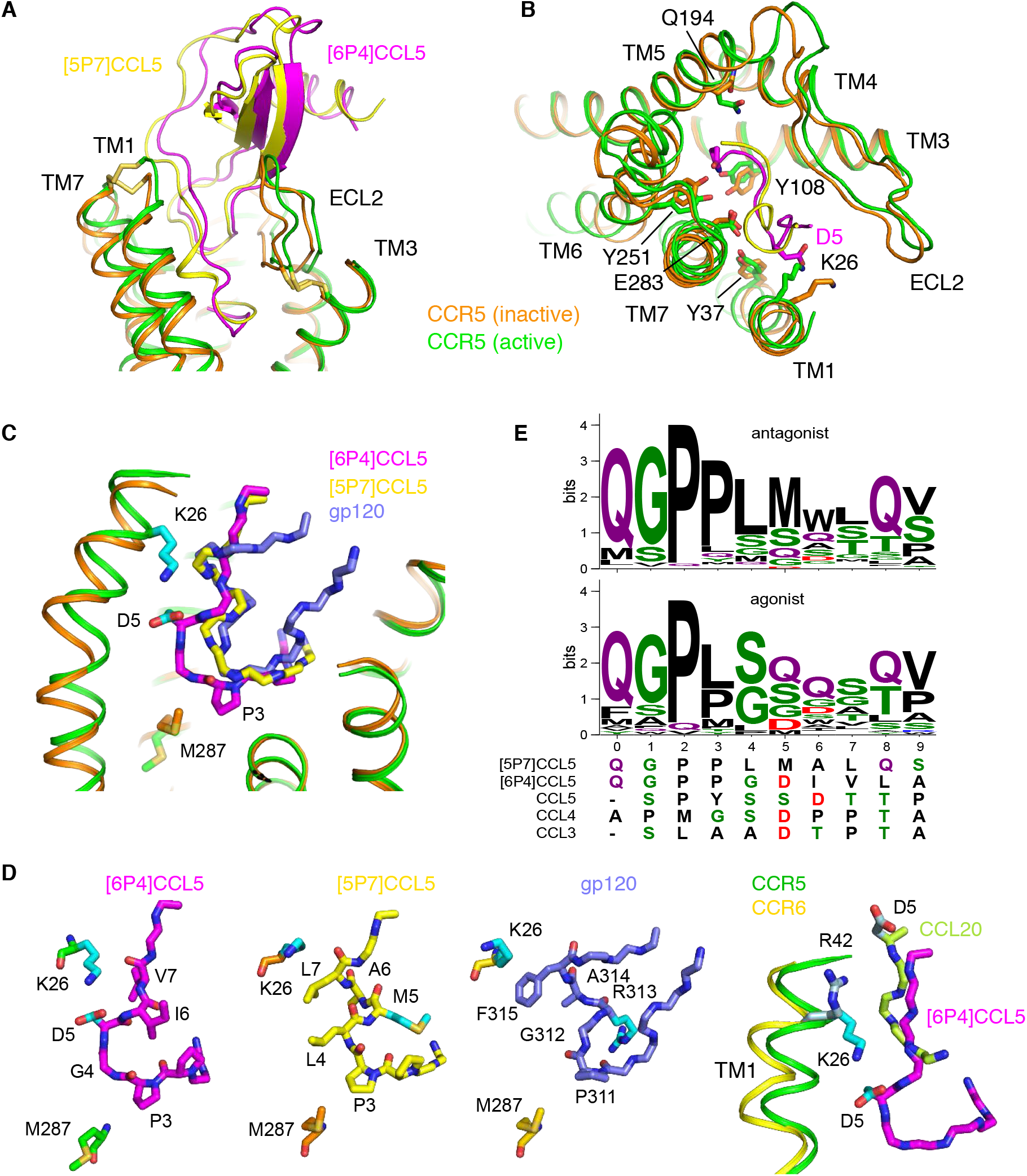
Activation mechanism of CCR5 by [6P4]CCL5 at CRS2. (A, B) Comparison between the insertion of the agonist [6P4]CCL5 (magenta) into active CCR5 (green) and the antagonist [5P7]CCL5 (yellow) into inactive CCR5 (orange; PDB ID: 5UIW) (A: side view; B: top view). Only the CCL5 N-terminal residues are shown in panel B. Important residues participating in the CCL5-CCR5 interaction are marked. (C) Comparison of insertion depths of agonist [6P4]CCL5 (magenta), antagonist [5P7]CCL5 (yellow), and the antagonist V3-loop of gp120 (slate; PDB 6MEO) into active CCR5 (green) and inactive CCR5 (orange). Salt bridge residues K26^1.28^-CCR5 and D5-[6P4]CCL5 (both in cyan) as well P3-[6P4]CCL5 and M287^7.43^ (in active and inactive CCR5) are shown as sticks. (D) Detailed view of [6P4]CCL5 N-terminus inserted into active CCR5, [5P7]CCL5 N-terminus, and gp120 V3-loop inserted into inactive CCR5; and comparison of the insertion of [6P4]CCL5 N-terminus into active CCR5 and of CCL20 N-terminus into active CCR6 (PDB ID: 6WWZ). (E) Sequence composition of the N-termini of CCL5 natural amino acid variants (Supplementary Table S2) with low (≤20%, N=83, top) and high (≥50%, N=34, bottom) Ca-signaling. The N-termini of [5P7]CCL5, [6P4]CCL5, and wild-type CCL3-5 (all agonists) are shown below.

The open conformation of the intracellular part of the active CCR5 differs from all inactive CCR5 structures thereby enabling the binding of the G protein (Fig. 1C): TM6 is moved outwards from the heptahelical bundle accompanied by further rearrangements of TM5, TM7, and intracellular loop 4 (ICL4). The moderate outward displacement of TM6 and the arrangement of G_i_ relative to CCR5 (Fig. 1A-C) agrees with previous GPCR•G_i_ complexes(*21, 22*).

## CRS1 interactions

The interactions between [6P4]CCL5 and CCR5 can be separated into the three canonical chemokine recognition sites (CRS) 1, 1.5, and *2(13, 23*). CRS1 consists of the contacts of the chemokine core with the extracellular side of the receptor and is dominated by electrostatic interactions. The core of [6P4]CCL5 sits on top of a wide opening in the extracellular part of the CCR5 transmembrane bundle, which is shaped by two disulfide bridges (C101^3.25^-C178^ECL2^, conserved in Class A GPCRs, and C20^N-term^-C269^7.25^, specific of chemokine receptors; superscripts indicate the GPCRdb numbering scheme(*24*)) (Fig. 1C). The [6P4]CCL5 strand β1 makes extensive contacts with polar residues in ECL2, while the CCR5 N-terminus directs towards a shallow grove between the chemokine N-loop and 40s-loop forming further extensive ionic and polar interactions. Due to missing density, the model of the CCR5 N-terminus could only be built starting from residue A16. Clear interactions are visible between CCR5 residues S17 and E18 and the chemokine residues R47 and Q48. To gain insights into the missing N-terminal region, CCR5 residues 1-15 were modelled (Fig. 1D) based on NMR NOE contacts(*25*) between wild-type CCL5 and an N-terminal fragment of CCR5 sulfated at residues Y10 and Y14. These post-translational modifications are important for chemokine affinity(*25–28*) and are expected to be present also in insect cell-expressed CCR5(*29*). The NMR structure of the CCR5-peptide•CCL5 complex(*25*) is completely compatible with the cryo-EM structure. The stability of the modeled interactions was assessed by molecular dynamics (MD) simulations of the CCR5/[6P4]CCL5 complex (Supplementary Video S2). The simulations reveal highly dynamic interactions between ^s^Y10 and ^s^Y14 of CCR5 and K45, R47 and R17 of [6P4]CCL5 in agreement with the NMR observations(*28*). In addition, a comparison of MD trajectories between non-sulfated and sulfated CCR5 indicates that sulfation induces a higher number of atomic contacts between the chemokine and the receptor N-terminus (Supplementary Fig. S5B), consistent with the higher affinity of the sulfated form(*27*).

## CRS2 interactions and activation

The N-terminus of [6P4]CCL5 reaches deep into the orthosteric pocket (CRS2) between the CCR5 7TM bundle. This contrasts with the shallow binding modes observed for the chemokines in the CCL20•CCR6•G_o_(*18*) and CXCL8•CXCR2•G_i_(*19*) complexes (see below). Of note, the N-terminal residues preceding C10 and C11 of monomeric CCL5 in solution have high nanosecond flexibility(*30*). However, they adopt a fixed conformation in the CCR5 complex. Residues 5-11 form the chemokine site (CS) CS1.5, which acts as a hinge between the chemokine core and CS2 (formed by residues 0-4 and located at the bottom of CRS2). Conspicuously, this very end of the [6P4]CCL5 N-terminus inserts several angstrom deeper into the CCR5 orthosteric pocket than [5P7]CCL5 or the V3-loop of gp120 in the respective inactive complexes with CCR5 (Fig. 2C), in a pose that partially overlaps with that of the antagonists maraviroc (Supplementary Fig. S6). Due to this deeper insertion, P3 of [6P4]CCL5 (but not maraviroc or P3 of [5P7]CCL5) can displace CCR5 M287^7.42^ and Y108^3.32^ apparently triggering the GPCR microswitch network (see below). The different insertion depth is caused by a markedly different structure of the chemokine hinge residues 5-11 (Fig. 2A-D) – a short helix in [5P7]CCL5 and an extended coil in [6P4]CCL5. This hinge is presumably the key for the placement of the chemokine N-terminus into CRS2, which controls receptor activation. In [6P4]CCL5, D5 of the extended hinge forms an ionic interaction with K26^1.28^ in TM1, whereas the side chain of the equivalent M5 of [5P7]CCL5 points in the opposite direction forming a helical turn. Apparently this helical turn is also pushed sideways by unfavorable interactions between [5P7]CCL5 L7 and K26^1.28^ (Fig. 2D). Very similar interactions and conformations are present in the inactive gp120•CCR5 complex, with F315, R313, and P311 taking the roles of [5P7]CCL5 L7, M5, and P3, respectively.

The structural finding that the CCL5 CS1.5 hinge conformation controls the insertion depth of residues 0-4 and thereby the activation state of CCR5 is corroborated by a statistical analysis of the pharmacological properties of CCL5 N-terminal amino acid variants. Currently, ~140 of these have been characterized for Ca-signaling (Supplementary Table S2), CCR5 internalization, and anti-HIV activity(*11*). A sequence analysis shows that residues 0-3 (highest abundance: QGPL) and 8, 9 (highest abundance: QV) are highly similar between 83 low- and 34 high-Ca-signaling N-terminal variants (Fig. 2E). The latter is not surprising since the selection libraries were to some extent biased towards these residues(*11*). In contrast, strong differences are observed for residues 4-7: for agonist variants the small, hydrophilic or negatively charged amino acids S, Q, G, D dominate, whereas antagonist variants contain mostly the large hydrophobic amino acids L, M, W, L. Apparently, the small hydrophilic residues direct the hinge towards K26^1.28^ in TM1, whereas the large hydrophobic residues make the hinge collapse to a helical turn. In agreement with their agonist pharmacology, both [6P4]CCL5 and wild-type CCL5 as well as the other major CCR5 agonist chemokines CCL3 (MIP1α) and CCL4 (MIP1β) contain an aspartic acid at positions 5 or 6 (Fig. 2E), which strengthens the extended hinge by forming a salt bridge to K26^1.28^.

As compared to [5P7]CCL5, the deeper binding pose of [6P4]CCL5 relocates its N-terminal pyroglutamate (PCA) group close to Y251^6.51^ and Q194^5.38^ (Fig. 2B) and our molecular dynamics simulations show that these groups interact through water-mediated hydrogen bonds (Supplementary Fig. S7). The placement of residue P3 of [6P4]CCL5 (Fig. 3) forces the relocation of M287^7.43^ in the receptor, which is accompanied by noticeable local changes in the backbone of TM7 (Supplementary Fig. S8) that bring the intracellular half of this helix towards the receptor core. This movement allows H289^7.45^ to push on W248^6.48^, possibly assisting the relocation of TM6 (Fig. 3A). P3 also lays on top of an ‘aromatic connector’ formed by CCR5 residues Y108^3.32^, F109^3.33^, F112^3.36^, and Y251^6.51^ (Fig. 3B), forcing the movement of Y108^3.32^ and resulting in a cascade of aromatic side chain relocations that transmit the activation signal to the receptor core. This allows the switch of the PIF motif (P206^5.50^, I116^3.40^, and Y244^6.44^) to an active conformation (Fig. 3D) and the large-scale movement of TM6. The relocation of TM6 and TM7 coincides with local structural changes in the NPxxY motif (Fig. 3C) leading to the formation of the conserved water-mediated interaction between Y297^7.53^ and Y2 1 4^5.58,^(*37*) and the opening of the binding pocket for H5 of G_i_, which includes R126^3.50^ in the ‘open’ conformation of the intra-helical ionic lock of the DRY motif (Fig. 3E and Supplementary Figure S9).

**Figure 3.**
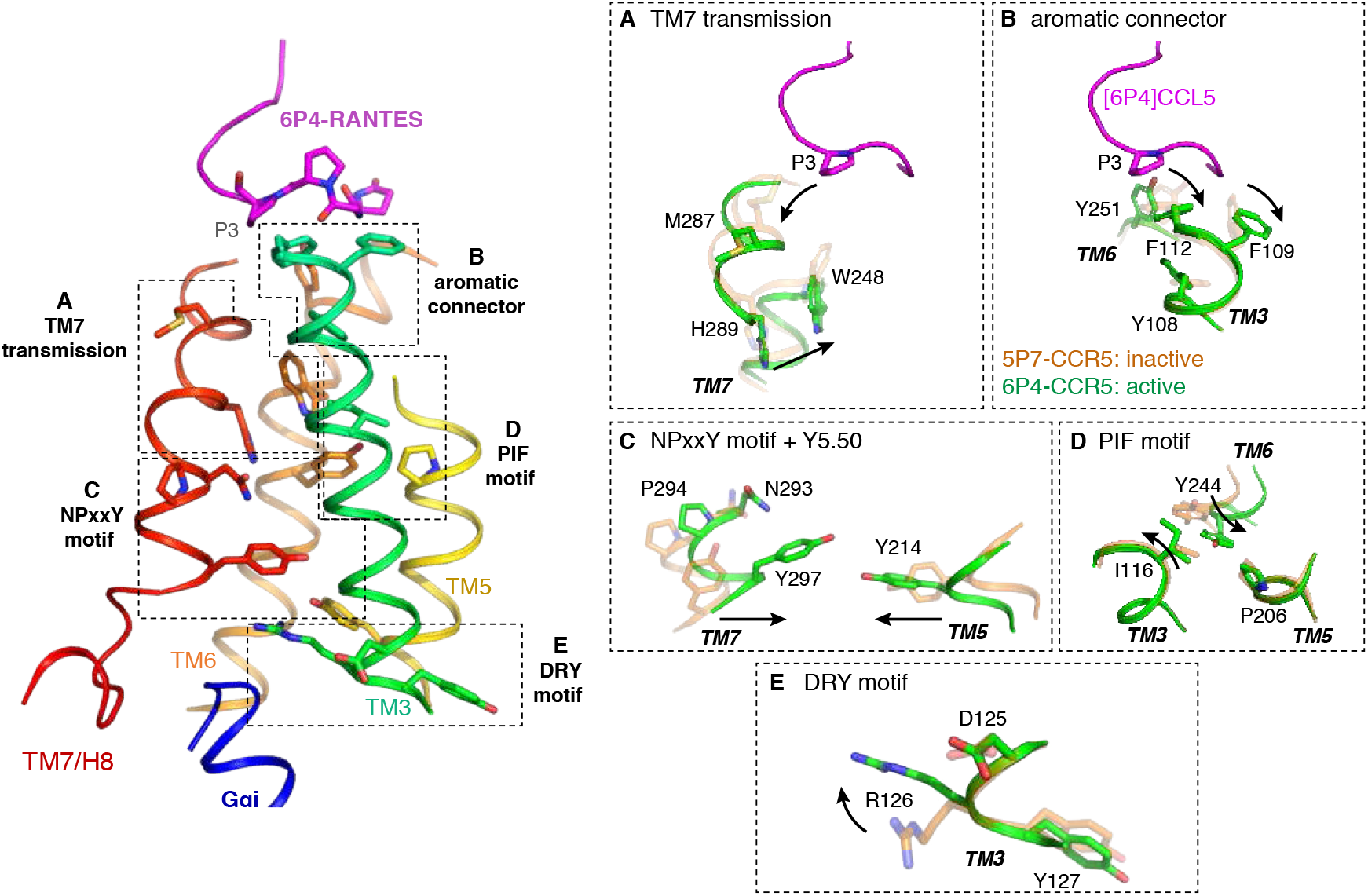
Transmission of the agonist signal to the receptor activation switches. Left: Activation switches in CCR5. Key residues are shown as sticks. For clarity, only part of the receptor structure is shown. (A) Transmission through TM7. (B) Transmission through the aromatic connector. (C) Activation of the NPxxY motif. (D) Activation of the PIF motif. (E) Activation of the DRY motif.

## Gi interactions

The binding interface of G_i_ to CCR5 is mediated exclusively by the Gα subunit (Supplementary Fig. S10) and can be divided into two main regions: the rim and the core (Fig. 4A and Supplementary Fig. S11). The rim contains two clearly separated parts (Fig. 4B): a ‘proximal’ side formed by the end of the αN helix (αNβ1) and nearby beta strands (β2β3) in Gα and ICL2 in the receptor, and a ‘distal’ side formed by beta strands (α4β6) in Gα and ICL3 in the receptor. In addition, the rim also includes interactions between α5 in G_i_ and ICL2/3 of the receptor. The core of the CCR5/Gi complex interface is formed exclusively by interactions of α5 in G_i_ (Fig. 4A,C). Here, the G_i_ α5 helix interacts with the cytoplasmic sides of TM2, TM3, and TM5 in one side of the core binding pocket, while the C-terminal hook of α5 (residues 352-354) leans towards TM6 and ICL4.

**Figure 4.**
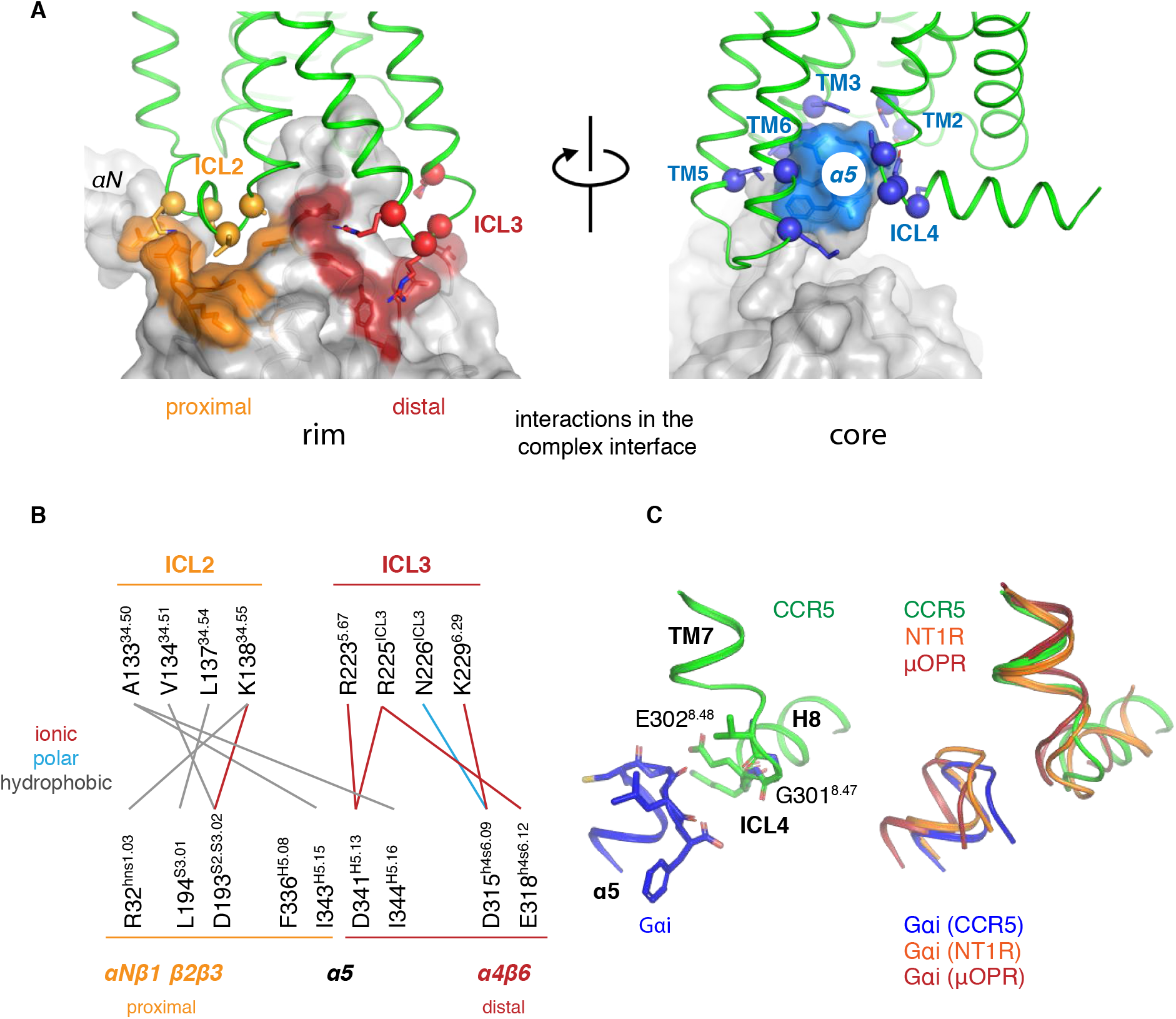
Binding interfaces between CCR5 and Giα. (A) Binding interfaces at the rim (left) and core (right) of the complex. Each interface is colored on the surface of the G protein and interacting residues in the receptor are shown as spheres (Cα) and sticks (side chains). (B) Residue-residue interactions at the rim of the binding interface. (C) Left: structure of the α5 hook and ICL4 in the [6P4]CCL5•CCR5 complex. Key residues are shown as sticks. Right: comparison between the α5 hook and ICL4 in the Gi-bound receptors CCR5, neurotensin type 1 (NT1R), and μ-opioid (μOPR).

These interfaces are common to all GPCR/G protein complexes, as they arise from the common overall relative orientation of the bound components. However, analysis of the currently available complexes reveals that the precise location and nature of the individual interface contacts vary to a certain degree (Supplementary Fig. S12). At the proximal rim of the interface, contacts are mostly hydrophobic and consistent with other G_i_ complexes. At the distal rim, we observe several ionic interactions absent in other structures. However, the most noticeable differences lie in the core region of the binding interface, where we observe different contacts between the hook of α5 (the last three C-terminal residues of G_i_) and ICL4 of CCR5. Interestingly, this is due to a distinct conformation of ICL4 of CCR5 in which G301^8.47^ and E302^8.48^ slightly relocate compared to other GPCR/Gi complexes, resulting in a different set of interactions between E302^8.48^ and the hook of α5 (Fig. 4B). The varying conformation of the short ICL4 illustrates the structural plasticity of this domain among Gi-binding GPCRs (Supplementary Fig. S13 and S14).

## Binding modes and activation mechanisms of long-vs. short-tail CC chemokines

The comparison between our structure and the inactive [5P7]CCL5•CCR5 complex(*13*) allows us to precisely pinpoint the activation mechanism of CCR5 by a chemokine agonist (Fig. 5A). The overall binding poses of the [5P7]CCL5 antagonist(*13*) and the [6P4]CCL5 agonist are similar, with the globular core of the chemokine held by the receptor N-terminus and ECL2 and the chemokine N-terminus reaching deep into the receptor transmembrane bundle. However, despite having the same 10-residue length, the N-termini of the two CCL5 derivatives differ in their amino acid sequences. This results in different chemokine/receptor interactions in this region: small, hydrophilic, or negatively charged residues in sequence positions 4-5 of [6P4]CCL5 lead to a straight hinge conformation that pushes residue P3 against the bottom of CRS2. In contrast, the large hydrophobic residues at these positions in [5P7]CCL5 force the hinge into a turn structure thereby making P3 recede. The more deeply inserted P3 of [6P4]CCL5 pushes onto the aromatic connector and residue M287^7.43^ thereby triggering the canonical activation switches resulting in the relocation of TM6/5/7 and the stabilization of the receptor active conformation. Notably, point mutations of the aromatic connector residues Y108^3.32^, F109^3.33^, and F112^3.36^ strongly reduce the efficacy of chemokine ligands while preserving their affinity(*32*). The highly conserved (~70% in non-olfactory human Class A GPCRs) residue W248^6.48^ lies at the center of these conformational changes, connecting the rearrangements at H289^7.45^ and Y244^6.44^ and, thus, the large-scale relocation of TM7 and TM6.

**Figure 5.**
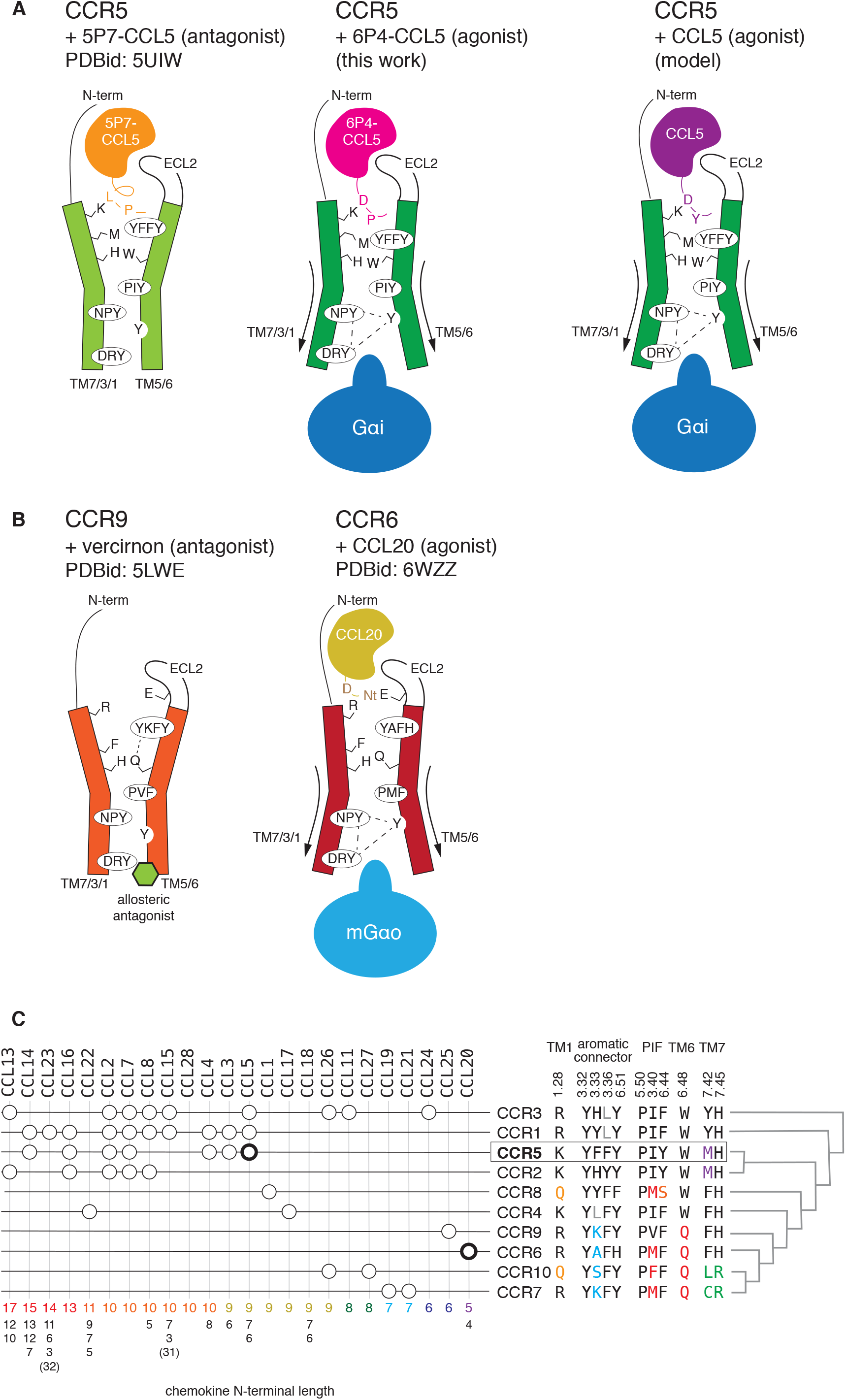
Activation mechanism of CCL chemokine receptors. (A) CCR5 bound to the antagonist [5P7]CCL5 (left; orange), the super-agonist [6P4]CCL5 (center; magenta), and the natural agonist CCL5 (right; purple). Key residues in the activation mechanism of CCR5 are shown. (B) Proposed activation mechanism for CCR6 by CCL20 (yellow; right) by comparing with the structure of CCR9 (left). (C) Pairing between CCR chemokine receptors and CCL chemokines(49). At the right, the sequence composition of key positions is shown, together with the phylogenetic relationship between the receptors. The lengths of the CCL chemokine N-termini according to UniProt(50) are shown at the bottom. The available active CCR/CCL complex structures are shown in bold.

We expect that, based on their N-terminal sequence, other identified CCR5 agonist or antagonist chemokines feature respective similar deeper (6P4[CCL5]-active-like) or less deep (5P7[CCL5]-inactive-like) positions of their N-terminal ends in CRS2. Also, using our structure as a template, we modeled the wild-type agonist CCL5 bound to CCR5 (Supplementary Fig. S15 and Supplementary Video S3). CCL5 – as the [6P4]CCL5 agonist – features an aspartate in its N-terminus (D6) able to interact with K26^1.28^. In CCL5, Y3 could be playing the role of P3 in [6P4]CCL5 by engaging the aromatic connector and M287^7.42^ (Fig. 5A). In the related chemokines CCL3 and CCL4, the aspartates at position 5 and the bulky residues at position 2 can carry out analogous functions.

The activation mechanism in CCL5/CCR5, in which the N-terminus of the chemokine reaches deep into the transmembrane bundle, differs substantially from that of CCL20/CCR6, where a much shorter CCL20 adopts a shallower binding pose and engages a non-canonical activation mechanism(*18*) (Fig. 2D, 5B). Thus, CC chemokine receptors can apparently be activated through two very different mechanisms by ‘long’ and ‘short’ chemokines. But what are the molecular features in the receptor that determine the type of activation? A phylogenetic analysis of CCR chemokine receptors (Supplementary Fig. S16) puts CCR5 and CCR6 into distinct distant subgroups. A more detailed sequence comparison of key residues in the activation mechanism shows that CCR chemokine receptors can be divided into two main groups according to the nature of the residue at position 6.48 (W vs Q) and, to some extent, of the aromatic connector (Fig. 5C). Remarkably, CC chemokine receptors featuring the conserved W^6.48^ (CCR1, CCR2, CCR3, CCR4, CCR5, and CCR8) tend to be more promiscuous and preferentially recognize chemokines with longer N-termini (9-14 residues) (Fig. 5C). On the other hand, CC chemokine receptors featuring Q^6.48^ (CCR6, CCR9, CCR7, and CCR10) bind to only a few (1-2) chemokines with short N-termini (4-9 residues). Interestingly, although position 6.48 allows for a certain degree of variability in human Class A GPCRs (70% W; 15% F; 5% Y; 10% other), a Q at this position is exclusive of this subgroup of chemokine human receptors, supporting the uniqueness of this ‘shallow’ activation mechanism.

## Conclusion

The structure of CCR5 in an active conformation allows us to elucidate a novel activation pathway of CC chemokine receptors by a chemokine agonist. In CCR5 and related receptors (CCR1, CCR2, CCR3, and CCR4), the respective cognate chemokines have long N-termini and bind deep into the orthosteric pocket (CRS2) thereby triggering the rearrangement of an aromatic connector in TM3 and TM6 and of the TM7 backbone. W^6.48^ lies at the center of these conformational changes connecting the receptor activation pathways through TM7 and TM6. In contrast, a subgroup of CC chemokine receptors (CCR6, CCR7, CCR9, and CCR10) harbor a Q residue at this position, a unique feature in human Class A GPCRs. The cognate chemokines of these receptors have shorter N-termini featuring a shallow binding mode and a specialized mode of activation. We expect that our findings will help to rationalize the relationship between sequence, structure, and activity of chemokines and their receptors and aid drug discovery.

## Methods

### Protein expression and purification

The wild-type human CCR5 gene containing a C-terminal 3C cleavage site followed by a FLAG-tag was cloned into the pFastBac1 vector and expressed in *Spodoptera frugiperda Sf9* insect cells using the baculoviral infection system. CCR5 expression and membrane preparation were performed as described(*15*). Membranes from 1-L culture of Sf9 cells were resuspended in 10 ml lysis buffer containing 2 mg/ml iodoacetamide, and EDTA-free complete protease inhibitor cocktail tablets, and incubated at 4°C for 1 hour. Then membranes were solubilized by supplementing 0.5 % LMNG at 4°C for 3 hours. The soluble fraction was isolated by centrifugation at 140,000 g and incubated with 1 ml of M2 Anti-FLAG affinity resin overnight at 4°C. The latter column was washed with 10 column volumes (CV) of washing buffer 1 (25 mM Hepes, 400 mM NaCl, 10% glycerol, 0.1% LMNG (w/v), pH 7.5), followed by 10 CV of washing buffer 2 (25 mM Hepes, 400 mM NaCl, 2 mM ATP, 5 mM MgCl2, 10% glycerol, 0.1% LMNG, pH 7.5) and subsequently washed with another 6 CV of washing buffer 1. The receptor was eluted with 3 CV of elution buffer consisting of 25 mM Hepes, 400 mM NaCl, 0.01% LMNG, 200 μg/ml FLAG peptide (DYKDDDDK), pH 7.5.

The DNA construct of [5P14]CCL5 cloned into a pET32a vector was a generous gift of Prof. LiWang. The DNA sequence of [6P4]CCL5 was obtained by mutating [5P14]CCL5 using standard QuickChange PCR. [6P4]CCL5 with enterokinase-cleavable N-terminal a thioredoxin fusion and hexa-histidine tags was expressed in the *E. coli* BL21 (DE3) strain cultured in Lysogeny broth media. Protein production was induced with 1 mM isopropyl β-D-thiogalactopyranoside when OD600 reached 0.7-0.8. After induction cells were grown for 20 h at 22°C and then harvested by centrifugation. 10 g of the cell pellet was resuspended in 50 ml of resuspension buffer (50 mM Tris, 6 M guanidinium HCl, 200 mM NaCl, pH 8.0) and lysed using a French press. The supernatant was isolated by centrifugation at 27,000g for 1 h and applied to a 5 ml HisTrap column. The column was washed with 10 CV of resuspension buffer and eluted with 3 CV of 60 mM NaOAc, 200 mM NaCl, 6 M guanidinium HCl. 20 mM β-mercaptoethanol was added to the elution fraction and incubated for 1 hour. The denatured protein was added dropwise into 250 ml of folding buffer (550 mM L-arginine hydrochloride, 20 mM Tris, 200 mM NaCl, 1 mM EDTA, 1 mM reduced glutathione, 0.1 oxidized glutathione, pH 8.0) and incubated overnight at 4°C. The solution was concentrated (MWCO 10 kDa) and dialyzed in 20 mM Tris, 200 mM NaCl, 2 mM CaCl_2_, pH 8.0. To cleave the fusion tags, enterokinase (NEB) was added, and the solution was incubated for 24 hours at room temperature. The protein was separated from the fusion tag using an acetonitrile gradient on a C4 reversed phase chromatography column (Vydac, Hesperia, CA) and then lyophilized. The lyophylisate was resuspended in 25 mM phosphate buffer, pH 6.0. The N-terminal amino acid of [6P4]CCL5 glutamine (Q0) was cyclized at 37°C for 48 hours.

The human Gα_i_ subunit (Gα_i1_) with an N-terminal TEV protease-cleavable deca-histidine tag was expressed in the *E. coli* BL21 (DE3) strain and purified as described(*21*).

The transducin heterotrimer was isolated from the rod outer segment of bovine retina (W L Lawson Company) and Gβ_1_γ_1_ was separated from Gα_t_ with Blue Sepharose 6 Fast Flow (GE Healthcare) as described(*21*). The Gα_i1_β_1_γ_1_ heterotrimer (G_i_) was prepared by mixing equimolar amounts of Gα_i1_ and Gβ_1_γ_1_ and incubated at 4°C for 1 h shortly before use for CCR5-G_i_ complex formation.

Fab16 was produced by papain digestion of IgG16 as described(*21*).

### Formation of the [6P4]CCL5•CCR5•G_i_•Fab16 complex

Pooled fractions of CCR5 eluted from the anti-FLAG resin and a molar excess of G_i_ heterotrimer were mixed together and incubated for 30 minutes. Then an equimolar amount of [6P4]CCL5 together with 25 mU/ml apyrase were added and incubated for another 2 hours. The complex was mixed with molar excess (1:1.4) of Fab16 and further incubated for at least 1 h. The mixture of [6P4]CCL5•CCR5•G_i_ and Fab16 was concentrated using an Amicon Ultra concentrator (MWCO 100 kDa) and loaded onto a Superdex 200 Increase 10/300 GL column for size-exclusion chromatography (SEC) with buffer containing 25 mM HEPES, 150 mM NaCl, 0.01% LMNG, pH 7.5. The protein quality of each fraction was evaluated by SDS-PAGE (Supplementary Fig. S1A,B). Fractions showing good purity and complex integrity were pooled together and concentrated for EM grid preparation.

### Cryo-EM sample preparation and image acquisition

For cryo-EM, 3.5 μL at 2.5 mg/ml sample was directly applied to glow-discharged 200 mesh carbon grids (Quantifoil R1.2/1.3). Grids were immediately plunge-frozen in liquid ethane using a FEI Vitrobot Mark IV (Thermo Fisher Scientific) with a blotting time of 3 s. The grids were screened for ice thickness and particle distribution using Glacios Cryo-TEM operated at 200 kV. Images were acquired from the selected grid using a Glacios Cryo-TEM (Thermo Fisher Scientific) operated at 200 kV equipped with a Gatan K3 Summit direct electron detector (Gatan Inc.). Automated data collection was carried out using SerialEM with a set of customized scripts enabling automated low-dose image acquisition(*33, 34*), and online pre-screened during data collection using FOCUS(*35*). Movie stacks of 40 frames were obtained with a defocus range of −1.0 to −2.0 μm at a magnification of 45000x (nominally 36000x) and the K3 detector operated in super-resolution mode (super-resolution pixel size, 0.556 Å). Each movie had a total accumulated dose exposure of ~49 e/Å^2^. A total of 2586 image stacks were collected for the [6P4]CCL5•CCR5•G_i_•Fab16 complex.

### Cryo-EM data processing

Contaminated micrographs were removed manually. Patch motion correction and Patch CTF parameter estimation were performed using algorithms implemented in CryoSparc v2.15.0(*36*). After sorting, micrographs with estimated resolution worse than 6.0 Å were discarded. The remaining motion-corrected summed images with dose-weighting were used for all other image processing in cryoSPARC. Approximately 2.6 million particles were auto-picked and subjected to several rounds of reference-free 2D classification to remove false positive particles. A total of 345,458 particles from 3D classes that demonstrated clear structural features were combined and subjected to 3D refinement, which led to a reconstruction at 3.6 Å resolution. Local refinement with a soft mask to exclude the density of the Gα α-helical domain and detergent micelle improved the overall resolution to 3.1 Å.

The final set of homogeneous [6P4]CCL5•CCR5•G_i_•Fab16 complex particles were subjected to 3D Variability Analysis (3DVA) implemented in CryoSPARC (https://www.biorxiv.org/content/10.1101/2020.04.08.032466v1.full.pdf). The solvent and micelle were excluded by applying a soft mask. The 3DVA was then run with this mask using three variability components and a low-pass filter resolution of 5 Å.

Reported resolutions calculated with a soft shape mask are based on the gold-standard Fourier Shell Correlation (FSC) using the 0.143 criterion. The local resolution was determined using ResMap(*37*).

### Model building and refinement

The crystal structures of the G_i_ heterotrimer (PDB ID: 5KDO), Fab16 (PDB ID: 6QNK and the [5P7]CCL5•CCR5 complex (PDB ID: 5UIW) were used as initial templates for model building. The models were docked into the 3D map as rigid bodies in Chimera(*38*). The [6P4]CCL5 N-terminus (up to the residue 8) was built *ab initio*. The remaining part of [6P4]CCL5 was taken from the 5UIW structure. Several rounds of manual building were performed in Coot(*39*). The model was finalized by refinement in Phenix 1.18.2.(*40*) against the 3.1 Å cryo-EM map. Structural figures were prepared in Chimera and PyMOL (https://pymol.org/2/). The refinement statistics are summarized in Supplementary Table S1.

### Amino acid sequence analysis

The analysis of N-terminal sequence similarity of the natural amino acid CCL5 variants (Supplementary Table S2) was carried using WebLogo(*41*).

### GTPγS test

The purified [6P4]CCL5•CCR5•G_i_ complex with or without Fab16 was incubated with 100 μM GTPγS in 25 mM HEPES, 150 mM NaCl, 0.01% LMNG, pH 7.5 for 1 h at 4°C followed by SEC analysis on a Superdex 200 Increase 10/300 monitoring the protein intrinsic tryptophan fluorescence with the excitation wavelength at 280 nm and emission wavelength 350 nm (Supplementary Fig. S1C).

### Homology modeling and molecular dynamics simulations

CCR5 N-terminal residues 1-25 were built with Modeller v9.16(*42*) using as templates i) residues 1-14 in the NMR solution structure of a doubly-sulfated (at Y10 and Y14) N-terminal segment of CCR5 bound to CCL5 (PDB ID: 6FGP) and ii) residues 25-320 in our [6P4]CCL5•CCR5•G_i_ cryo-EM model. The chemokines were used as a guide for the structural alignment of the templates. Secondary structure helical restraints were added to residues 21-24 of CCR5. Cysteine bridges in CCR5 (C20-C269^7.24^ and C101^3.25^-C178^45.50^) and in [6P4]CCL5 (C10-C34 and C11-50) were explicitly defined during model building. All models were subjected to 300 iterations of variable target function method optimization and thorough molecular dynamics and simulated annealing optimization and scored using the discrete optimized protein energy potential. The 20 best-scoring models were analyzed visually, and a suitable model (in terms of low score and overall structure) was selected (Figure 1D, right panel).

This model of CCR5 (residues 1-320) bound to [6P4]CCL5 was used for molecular dynamics simulations of the non-sulfated and sulfated (Y10 and Y14) forms. Coordinates were first pre-processed using VMD1.9.3(*43*). The receptor-ligand complex (i.e. CCR5-[6P4]CCL5 or CCR5-CCL5) was then embedded into a 90 Å x 90 Å lipid bilayer composed of 80% 1-palmitoyl-2-oleoyl-sn-glycero-3-phosphocholine (POPC) and 20% cholesterol. The system was solvated with explicit water molecules, neutralized, and its ionic strength was adjusted using the CHARMM-GUI builder(*44*). Disulfide bridges were explicitly defined between: C50-C11 and C34-C10 in CCL5 or [6P4]CCR5, and C101^3.25^-C178, and C20-C269^7.25^ in CCR5. Except for CCR5 residues D76^2.50^, E283^7.39^, and E302^8.48^, which were protonated, all titratable residues of CCR5 and CCL5 were left in their dominant protonation state at pH 7.0. Prior to production runs, the geometry of the system was optimized by energy minimization and further relaxed by a sequence of equilibration steps where harmonic positional restraints were applied to all Cα atoms of the protein, and gradually released throughout the equilibration. In the last equilibration step (i.e. before completely releasing all protein restraints), water, ion and lipids were allowed to diffuse without restraints during 50 ns to allow for adequate equilibration of the lipid mixture. Upon equilibration was completed, five independent trajectories of each system were spawned from the last snapshot of the equilibrated trajectory using a random seed. Production simulations for each replica were run in the NPT ensemble at 1,013 bar and 310 K for 500 ns each. All simulations were run using Gromacs v2020(*45*) with the CHARMM36m force field(*46*). Gromacs v2020 and VMD1.9.31, were used to post process and analyze all trajectories. MD simulation figures were rendered using VMD1.9.3, and the R ggplot2 library(*47*).

The equilibrated model [6P4]CCL5 bound to CCR5 was used to model the binding pose of the wild type CCL5. The sequence of CCL5 was threaded on [6P4]CCL5 (6P4: QGPPGDIVLACC / CCL5: SPYSSDTTP-CC) and steric clashes were relieved using the molecular graphics software PyMOL. Using this structure as a template, residues 1-9 of CCL5 and all residues 8 Ångstrom around Y3 of CCL5 were re-modeled with Modeller v9.16 using the protocol described above. The stability of the resulting binding pose was assessed by molecular dynamics simulations using the protocol described above.

A list of simulations performed in this work is given in Supplementary Table S3. Molecular dynamics simulations were performed at the Paul Scherrer Institute computing cluster and at the Swiss National Supercomputing Centre (CSCS).

Electrostatic potentials were calculated using the APBS method(*48*) as implemented in PyMOL using a concentration of 0.150 M for the +1 and 1 ion species. The biomolecular surface is colored from red (5 kT/e) to blue (+5 kT/e) according to the potential on the soluble accessible surface.

## Acknowledgments

This work was supported by the Swiss National Science Foundation (grants 149927 and 173089 to S.G.; grant 192780 to X.D.; SNF R’EQUIP 177084 to T.M.; NCCR TransCure to H.S.; SNF Sinergia 183563 to G.F.X.S.), the Synapsis Foundation (grant 2018-P104 to G.F.X.S.), and the European Union (grants FP6-EMPRO and FP7-CHAARM to S.G.). We gratefully acknowledge Mohamed Chami and Lubomir Kovacik (Biozentrum BioEM Lab) for help with cryo-EM data collection, Patricia LiWang (University of California) for a generous gift of the [5P14]CCL5 DNA construct, Marco Rogowski (Biozentrum) for preparation of [6P4]CCL5, Jonas Muehle (Paul Scherrer Institute) for preparation of Fab16, Marc Caubet (High Performance Computing and Emerging technologies Group, Paul Scherrer Institute) for technical support with molecular dynamics simulations, Shin Isogai for helpful discussion, as well as the sciCORE facility of the University of Basel for the computer infrastructure.

## Data availability

The cryo-EM map of [6P4]CCL5•CCR5•G_i_•Fab16 has been deposited in the Electron Microscopy Data Bank as entry EMD-XXX and the corresponding model in the Protein Data Bank as entry XXX.

## Author contributions

P.I., A.G. and S.G. conceived the study. P.I., C.-J.T. and F.P. expressed and purified proteins and developed the protocol for forming the [6P4]CCL5•CCR5•G_i_•Fab16 complex. A.G. assisted with the expression and purification of CCR5. P.I. and C.-J.T. prepared cryo-EM grids. K.G. and P.I. collected the cryo-EM data. P.I. and N.D. processed the cryo-EM data, built and refined the model. X.D. and R.G designed and performed molecular dynamics simulations. H.S and T.M. provided guidance on EM sample preparation, data collection, and model refinement. P.I., X.D., O.H. and S.G. analyzed the structure and wrote the manuscript with input from C.-J.T., A.G. and G.S.

## List of Supplementary Materials

Table S1 – S3

Fig S1 – S16

Movies S1 – S3

