## Supplementary Figures for "Structural basis of the activation of the CC chemokine receptor 5 by a chemokine agonist"

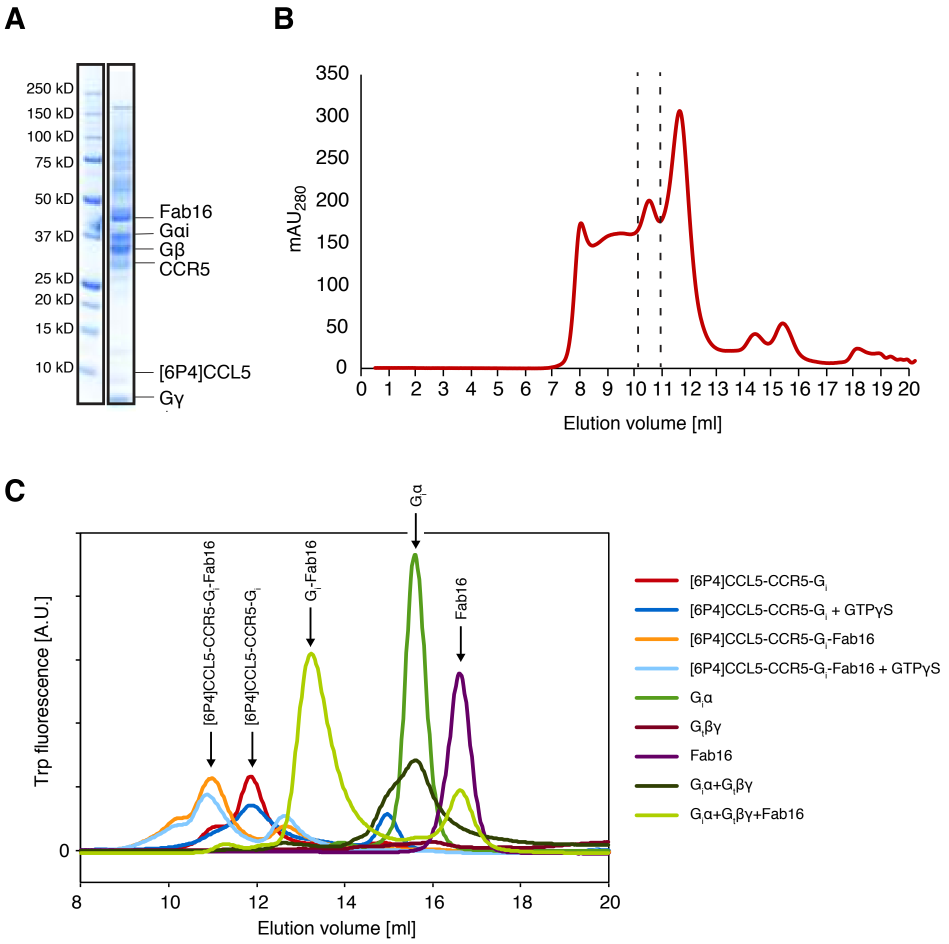


**Figure S1. Purification of the [6P4]CCL5•CCR5•$G_i$•Fab16 complex** (A) SDS-PAGE analysis and (B) Superdex200 size-exclusion chromatography elution profile of the [6P4]CCL5•CCR5•$G_i$•Fab16 complex. The dashed lines in B indicate the part of the elution volume used for cryo-EM sample preparation. (C) Analytical size-exclusion chromatography was used follow complex formation at different stages as well as the stability of the complexes in the presence and absence of Fab16 and GTPγS. The integrity of the [6P4]CCL5•CCR5-$G_i$ complex (red) is disturbed in the presence of GTPγS (dark blue). Addition of Fab16 (orange) stabilizes the complex in the presence of GTPγS (light blue), as the antibody fragment constrains the conformational flexibility of the $G_i$ heterotrimer.

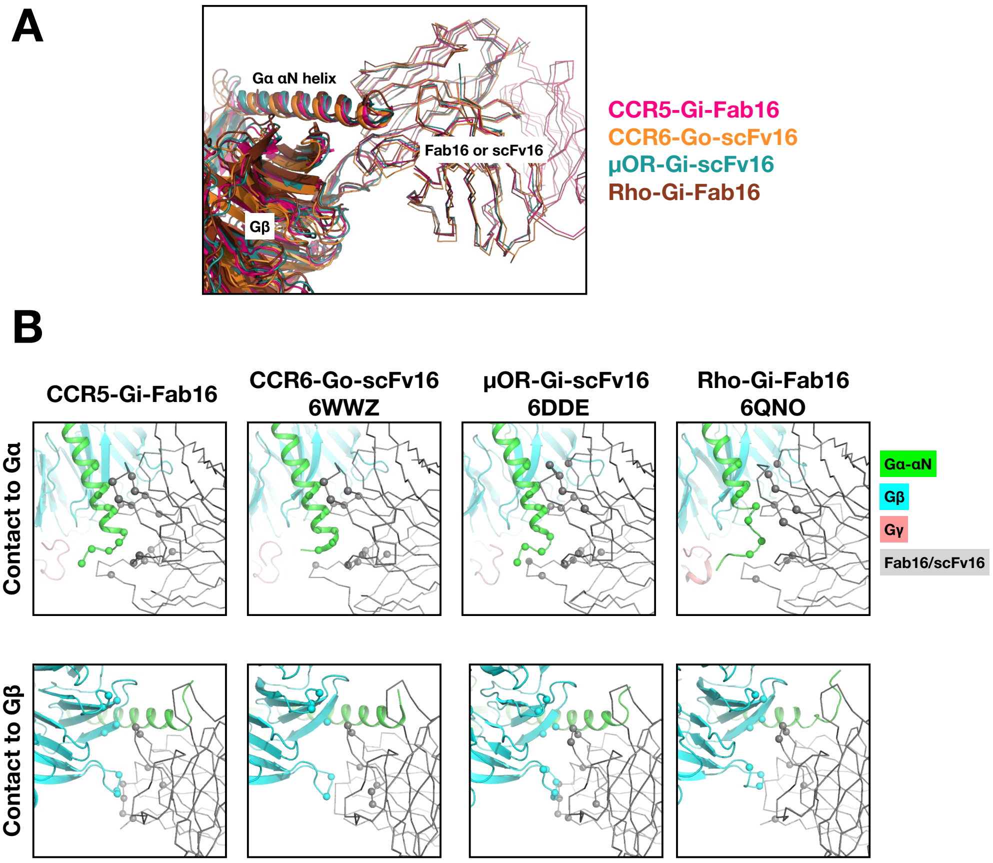


**C**

**Contact of Gα to Fab16/scFv16**

```
               4                  22
Giα-CCR5       TLSAEDKAAVERSKMIDRN
Goα-CCR6       -LSAEERAALERSKAIENK
Giα-uOR        -LSAEDKAAVERSKMIDRN
Giα-Rho        TLSAEDKAAVERSKMIDRN
```

**Contact of Gβ to Fab16/scFv16**

```
               66             81        91  // 110                        131    137
Gβ-CCR5        DSRLLVSASQDGKLIIWDSYTTNKVHAI // NYVACGGLDNICSIYNLKTREGNVRVSR
Gβ-CCR6        DSRLLVSASQDGKLIIWDSYTTNKVHAI // NYVACGGLDNICSIYNLKTREGNVRVSR
Gβ-uOR         DSRLLVSASQDGKLIIWDSYTTNKVHAI // NYVACGGLDNICSIYNLKTREGNVRVSR
Gβ-Rho         DSRLLVSASQDGKLIIWDSYTTNKVHAI // NYVACGGLDNICSIYNLKTREGNVRVSR
```

**Figure S2. Contacts between Fab16 or scFv16 and Gα/β in existing GPCR/$G_{i/o}$ protein complexes.** (A) Structural superposition of GPCR-$G_{i/o}$ complex structures at the interface between G protein α/β subunits and Fab16 or scFv16. Structures of CCR5-$G_i$ (this structure), CCR6-$G_o$ (PDB ID: 6WWZ), μ-opioid receptor-$G_i$ (PDB ID: 6DDE), and rhodopsin-$G_i$ (PDB ID: 6QNO) are aligned to the scFv16 chain. (B) Detail of the contact interfaces between Fab16 or scFv16 and the αN helix of Gα (top) and Gβ (bottom). Contact residues were selected within 4 Å of the neighboring molecule and are shown as spheres (only Cα). (C) Sequence alignment of the Gα and Gβ residues forming contacts with the antibody fragments. Residues of Gα (top panel) and Gβ (bottom panel) within 4 Å of Fab16 or scFv16 are highlighted in green and cyan, respectively. The comparison shows how the binding interfaces are very similar in all complexes.

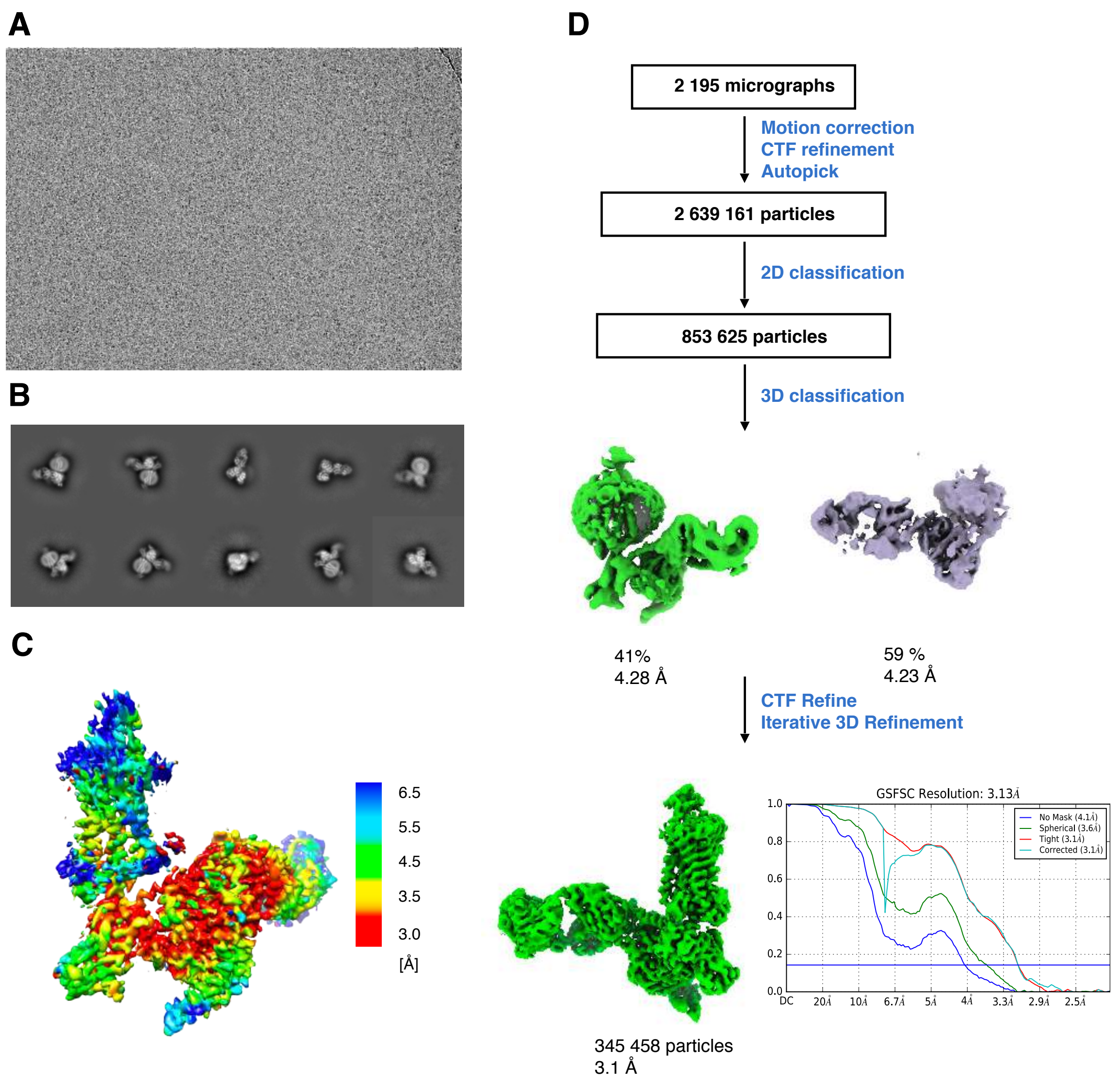


**Figure S3. Cryo-EM data processing.** (A) Representative cryo-EM micrograph of the [6P4]CCL5•CCR5•$G_i$•Fab16 complex sample and (B) representative 2D class averages showing distinct views and structural features. (C) Density map colored by local resolution. (D) Workflow of cryo-EM data processing. The Fourier shell correlation (FSC) curve indicates an overall nominal resolution of 3.1 Å using the Gold-standard FSC=0.143 criterion.

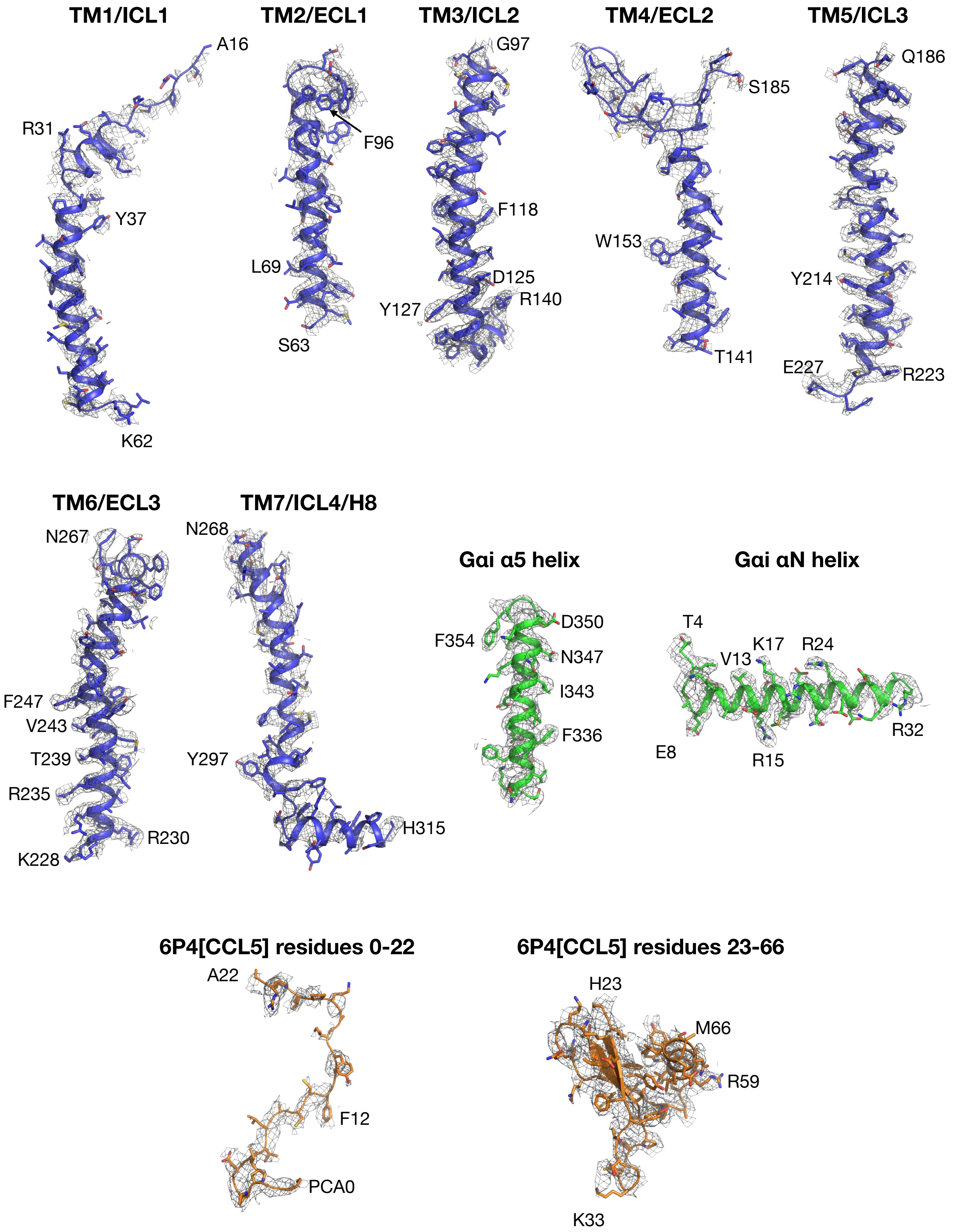


**Figure S4. Atomic model of [6P4]CCL5•CCR5•Gi-Fab16 in the cryo-EM density map.** Transmembrane helices TM1-TM7/H8 of CCR5 (blue cartoons) are shown together with the cryo-EM density map displayed at a 7.5 σ cut-off within 2 Å of the model. The αN and α5 helices of Giα1 (green), and [6P4]CCL5 residues 0-22 and 23-66 (orange) are shown in the same representation.

A Conformational dynamics of 6P4[CCL5] in molecular dynamics simulations.

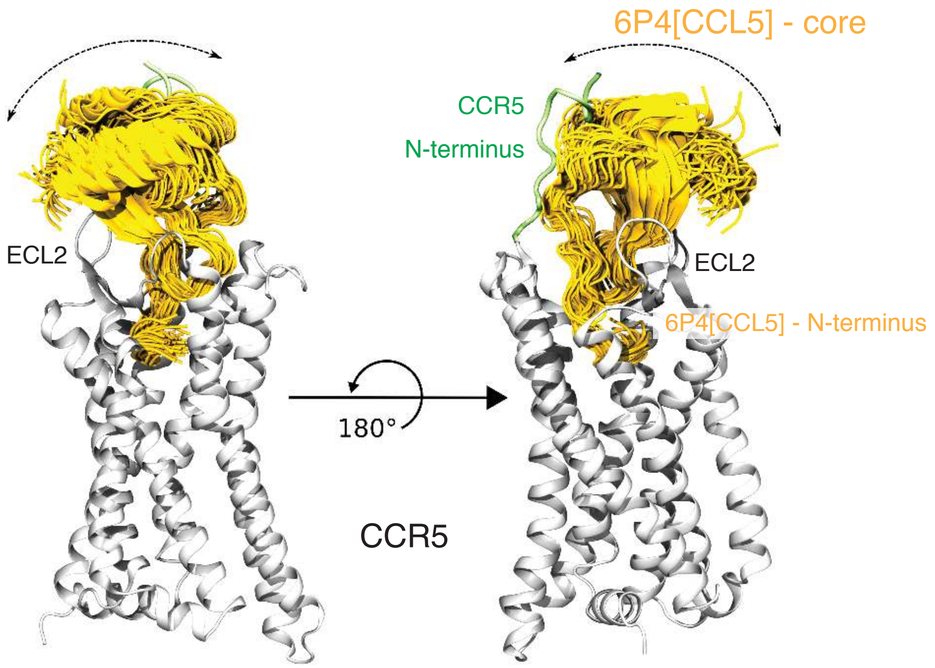


B Number of contacts between the N-terminus of CCR5 and 6P4[CCL5] in molecular dynamics simulations. TYR = non-sulfated Y10 and Y14. TYS = sulfated Y10 and Y14.

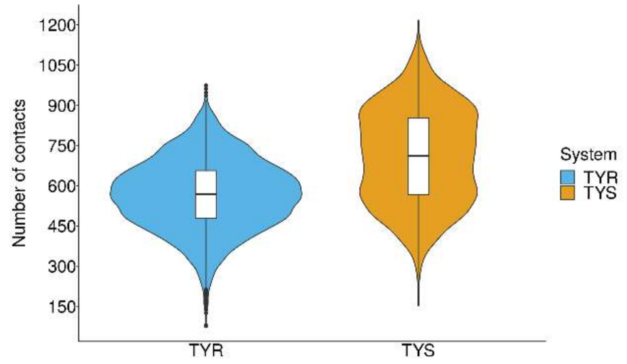


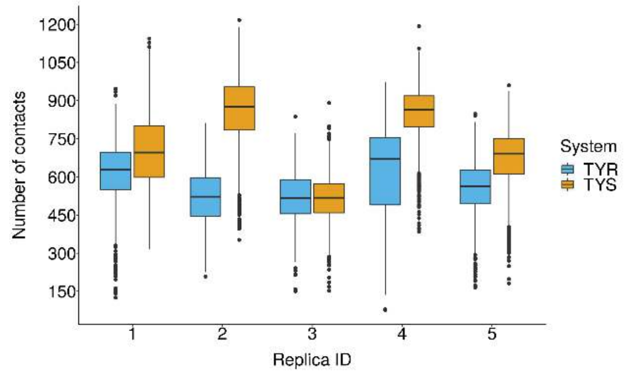


**Figure S5. Effect of tyrosine sulfation on the interaction between the CCR5 N-terminus and CCL5.** (A) Representative conformational ensemble of [6P4]CCL5 – with sulfated Y10 and Y14 – in MD simulations. In each frame, the structure of the N-terminal residues (0-10) of the chemokine are superposed. Only one receptor is shown for clarity. The core domain of the chemokine (bound between the N-terminus and extracellular loop 2 (ECL2) of the receptor) is relatively flexible. (B) Number of contacts per frame established between the CCR5 N-terminus (residues 8 to 18) and [6P4]CCL5 in MD simulations of the absence (TYR) and presence (TYS) of sulfation at residues Tyr10 and Tyr14 of CCR5 N-terminus. The top panel shows the number of contacts per frame across the accumulated simulation time for each system (i.e. 5 x 500 ns = 2.5 μs). The bottom panel shows the number of contacts per frame in each independent simulation replica of 500 ns. In the simulations, tyrosine sulfation results in a higher number of contacts between the receptor and the chemokine.

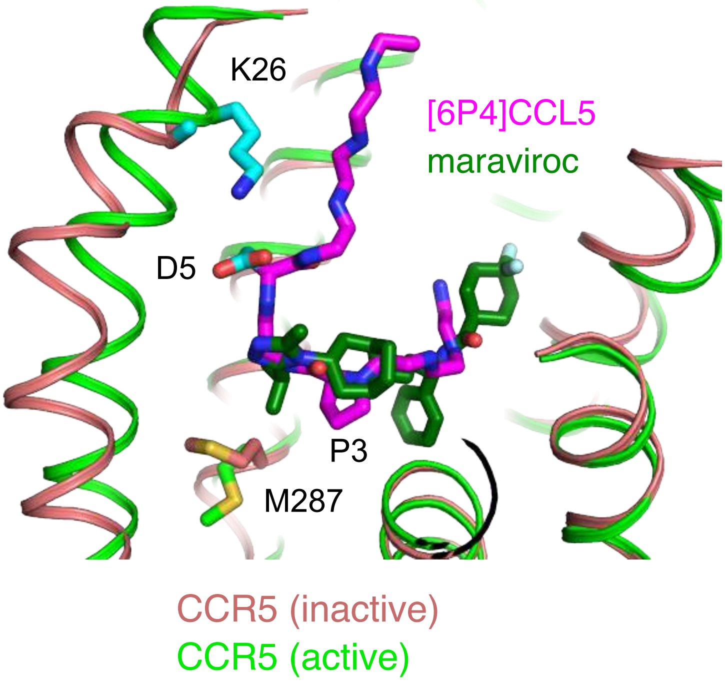


**Figure S6. Comparison between the binding poses of the agonist [6P4]CCL5 N-terminus and the antagonist maraviroc in CRS2 of CCR5.** Same view of CRS2 as Fig. 2C, showing the agonist [6P4]CCL5 (magenta) bound to active CCR5 (green; this structure) and the antagonist maraviroc (dark green) bound to inactive CCR5 (brown-orange; PDB ID: 4MBS). The salt bridge residues K26$^{1.28}$-CCR5 and D5-[6P4]CCL5 (both in cyan) as well P3-[6P4]CCL5 and M287$^{7.43}$ (in active and inactive CCR5) are shown as sticks. The azabicyclo group of maraviroc does not insert as deeply as P3 of [6P4]CCL5 at the equivalent position. As a consequence, the sidechain of M287$^{7.43}$ remains in the inactive conformation (see Fig. 2C). The phenyl group of maraviroc, instead, reaches deeply into the receptor between the residues of the aromatic cluster, possibly blocking its function as a signal relay.

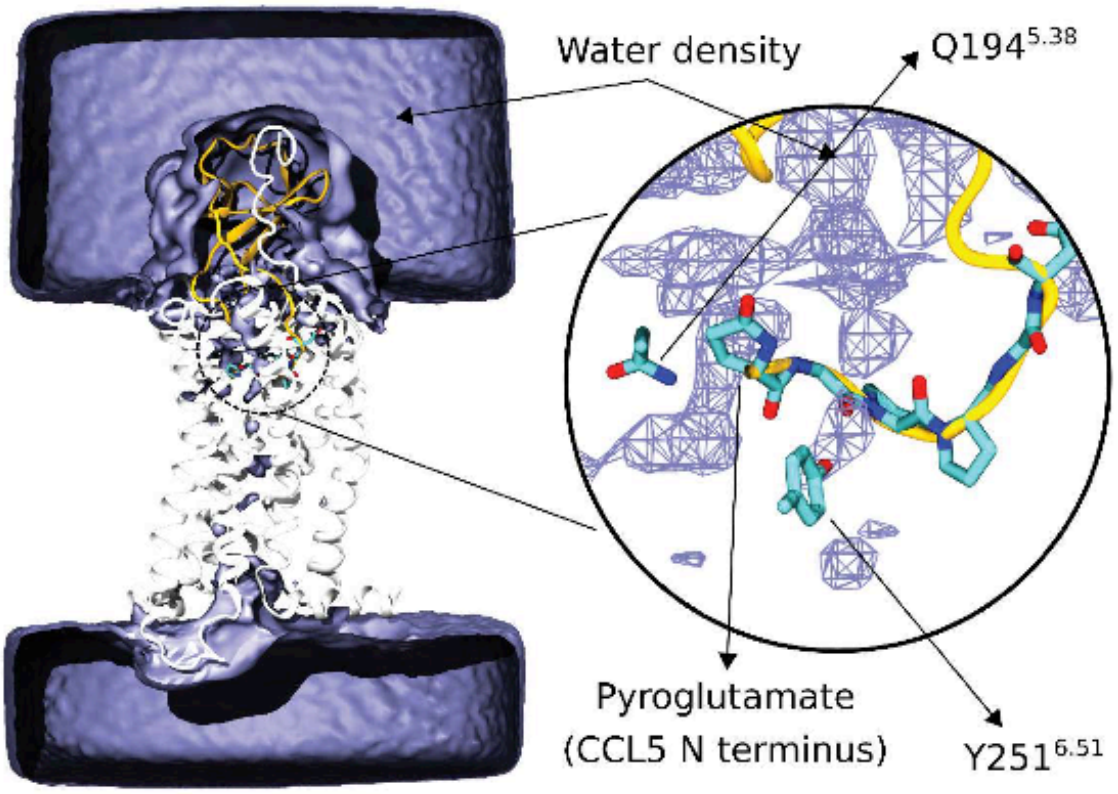


**Figure S7. Water density in molecular dynamics simulations of the CCR5/6P4[CCL5] complex.** All replicas (see Supplementary Table S3) were used to compute the average water density (blue surface). The inset shows a closer view of the average water density (blue mesh) solvating the N terminal pyroglutamate group of [6P4]CCL5 (yellow cartoon).

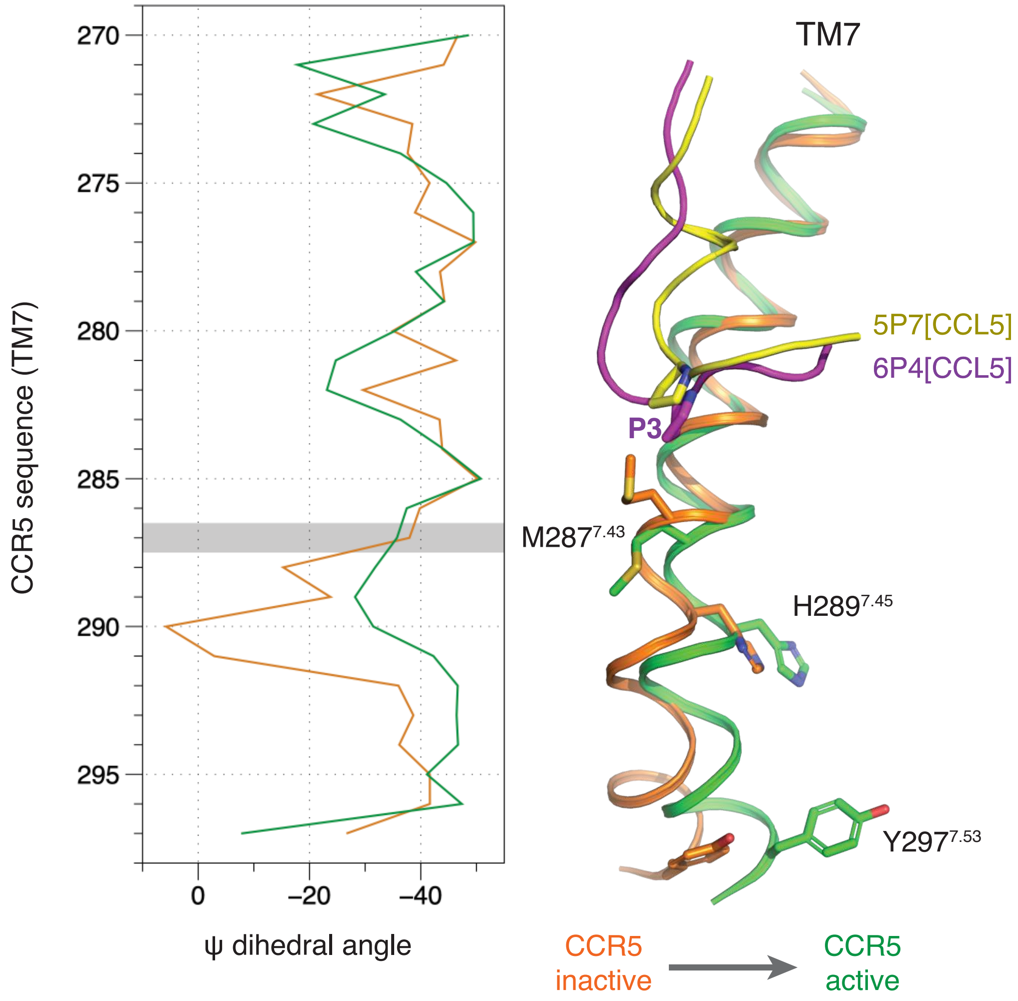


**Figure S8. Comparison of the structure of TM7 between inactive [5P7]CCL5-bound and active [6P4]CCL5-bound CCR5.** Left: ψ dihedral angles along TM7 reveals significant changes in the structure of the helical backbone at residues T288$^{7.44}$ – C291$^{7.47}$. Right: These changes occur together with the relocation of the M287$^{7.43}$ side chain (forced by the deeper insertion of the chemokine N-terminus at P3) and translate into the movement of the cytoplasmic side of TM7 (and the key residues H289$^{7.45}$ and Y297$^{7.53}$) towards the receptor core.

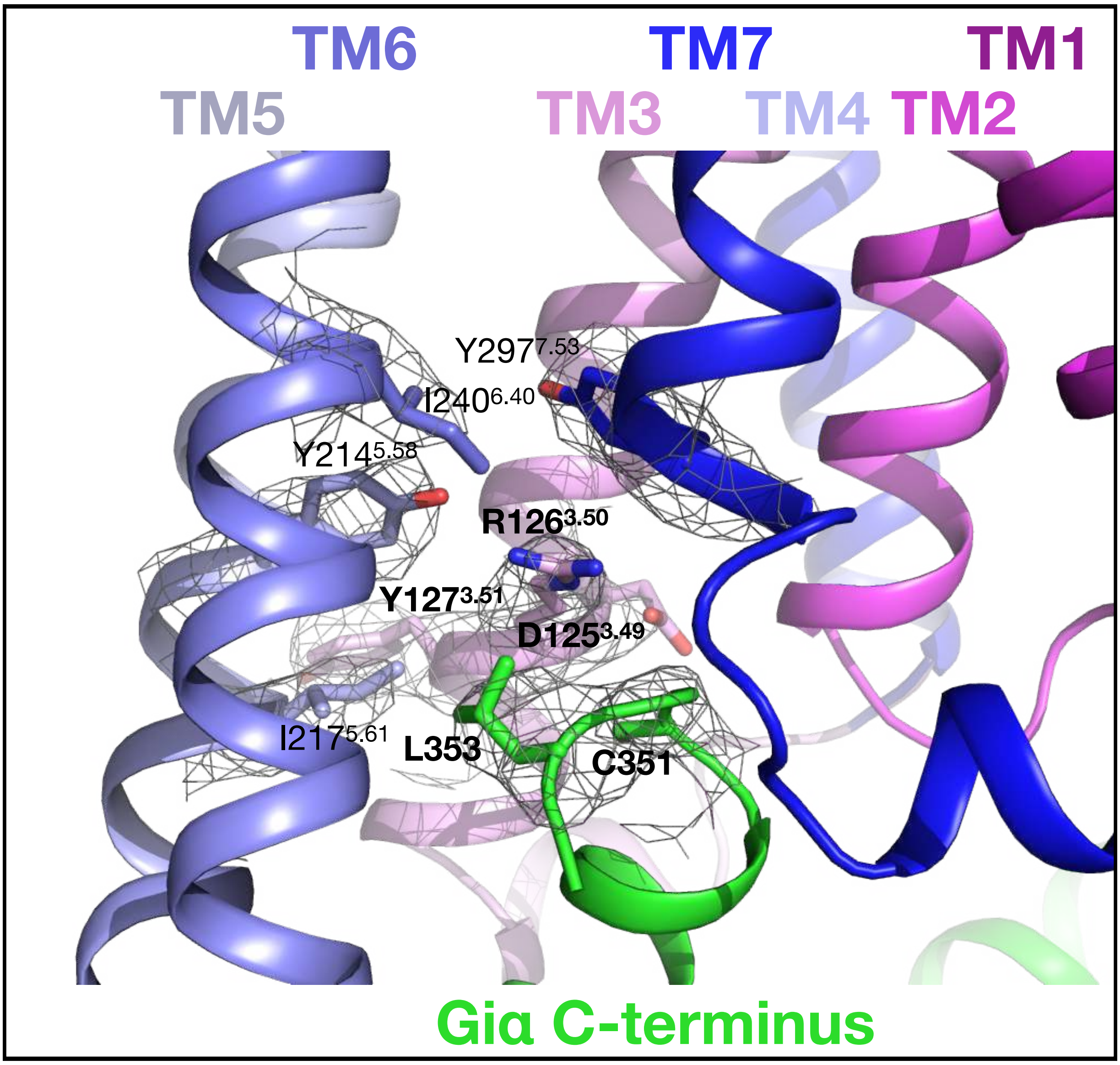


**Figure S9. Electron density around the receptor DRY motif in the [6P4]CCL5•CCR5•G$_i$-Fab16 structure.** The DRY motif (D125$^{3.49}$, R126$^{3.50}$, Y127$^{3.51}$) of CCR5 and residues within 4 Å of R126$^{3.50}$ are shown as sticks. The cryo-EM density around these residues is displayed at a 7.5 σ cut-off as a grey mesh. The opening of the ionic lock between D125$^{3.49}$ and R126$^{3.50}$ allows R126$^{3.50}$ to engage Y214$^{5.58}$ and Y297$^{7.53}$, leading to the formation of a cytoplasmic cleft that allows binding of the α5 helix in the Gα protein. R126$^{3.50}$ directly interacts with the α5 hook via C351$^{H5.23}$ and L353$^{H5.25}$ of Giα.

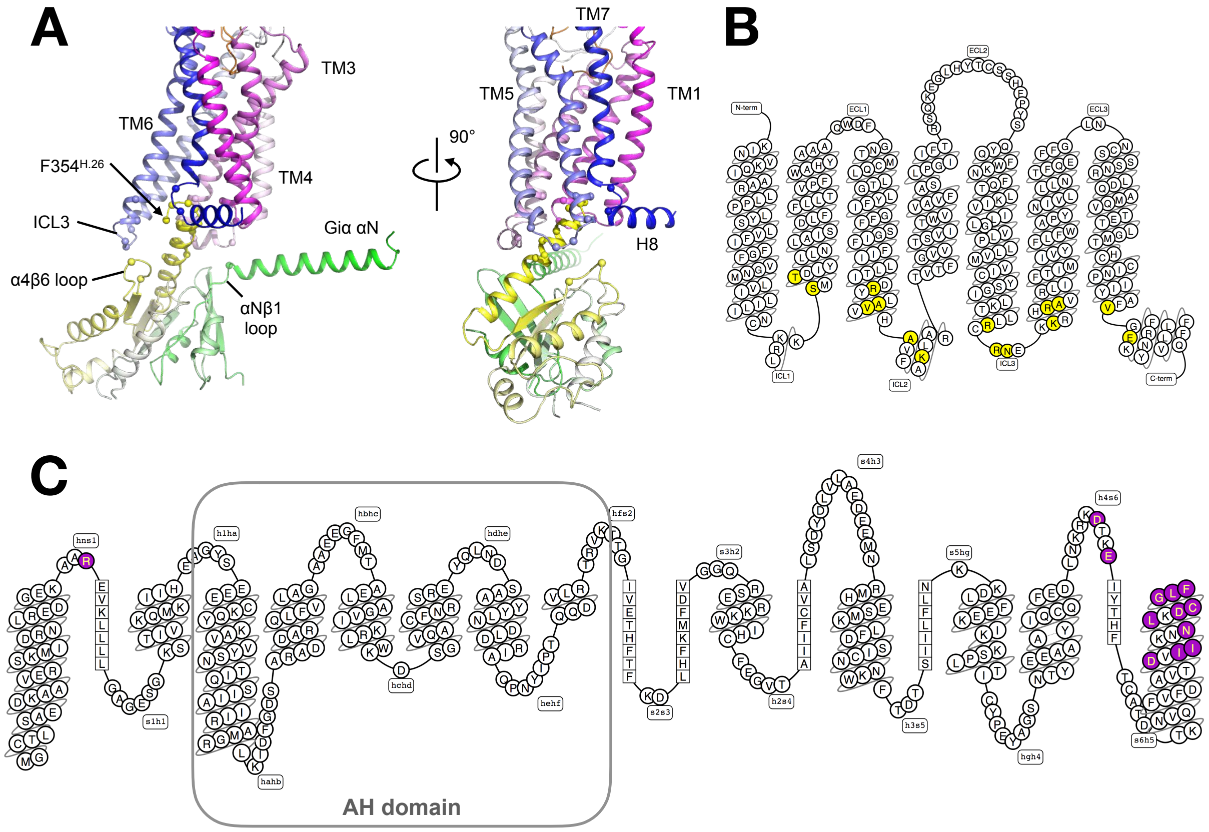


**D**

| Giα | αNβ1 | α4β6 | | α5 helix | | | | | | | | | |
|---|---|---|---|---|---|---|---|---|---|---|---|---|---|
| **CCR5** | **hns1.03 R32** | **h4s6.09 D315** | **h4s6.12 E318** | **H5.13 D341** | **H5.15 I343** | **H5.16 I344** | **H5.19 N347** | **H5.20 L348** | **H5.22 D350** | **H5.23 C351** | **H5.24 G352** | **H5.25 L353** | **H5.26 F354** |
| **TM2** | | | | | | | | | S63$^{2.37}$ T65$^{2.39}$ | | | | |
| **TM3** | | | | | | V130$^{3.54}$ | A129$^{3.53}$ | V130$^{3.54}$ | | R126$^{3.50}$ | | R126$^{3.50}$ | |
| **ICL2** | K138$^{34.55}$ | | | | A133$^{34.50}$ | A133$^{34.50}$ | | | | | | | |
| **TM5** | | | | R223$^{5.67}$ | | R223$^{5.67}$ | | | | | | | |
| **ICL3** | | N226 | R225 | | | | | | | | | | |
| **TM6** | | | | | | | | | | | | R232$^{6.32}$ | K229$^{6.29}$ R232$^{6.32}$ A233$^{6.33}$ |
| **TM7** | | | | | | | | | | | V300$^{7.56}$ | | |
| **H8** | | | | | | | | | | | E302$^{8.48}$ | | |

**Figure S10. Contact interface between CCR5 and Giα.** (A) Structure of CCR5 and Giα in the complex colored in blue-white-magenta (receptor) and green-white-yellow (G protein) spectra. The α carbon atoms of contact residues (within 4 Å) are shown as spheres. (B) Snake plot of CCR5; contacts with Giα are colored in yellow. (C) Snake plot of Giα1; contacts with CCR5 are colored in purple. (D) Table of residue-residue contacts between CCR5 and Giα.

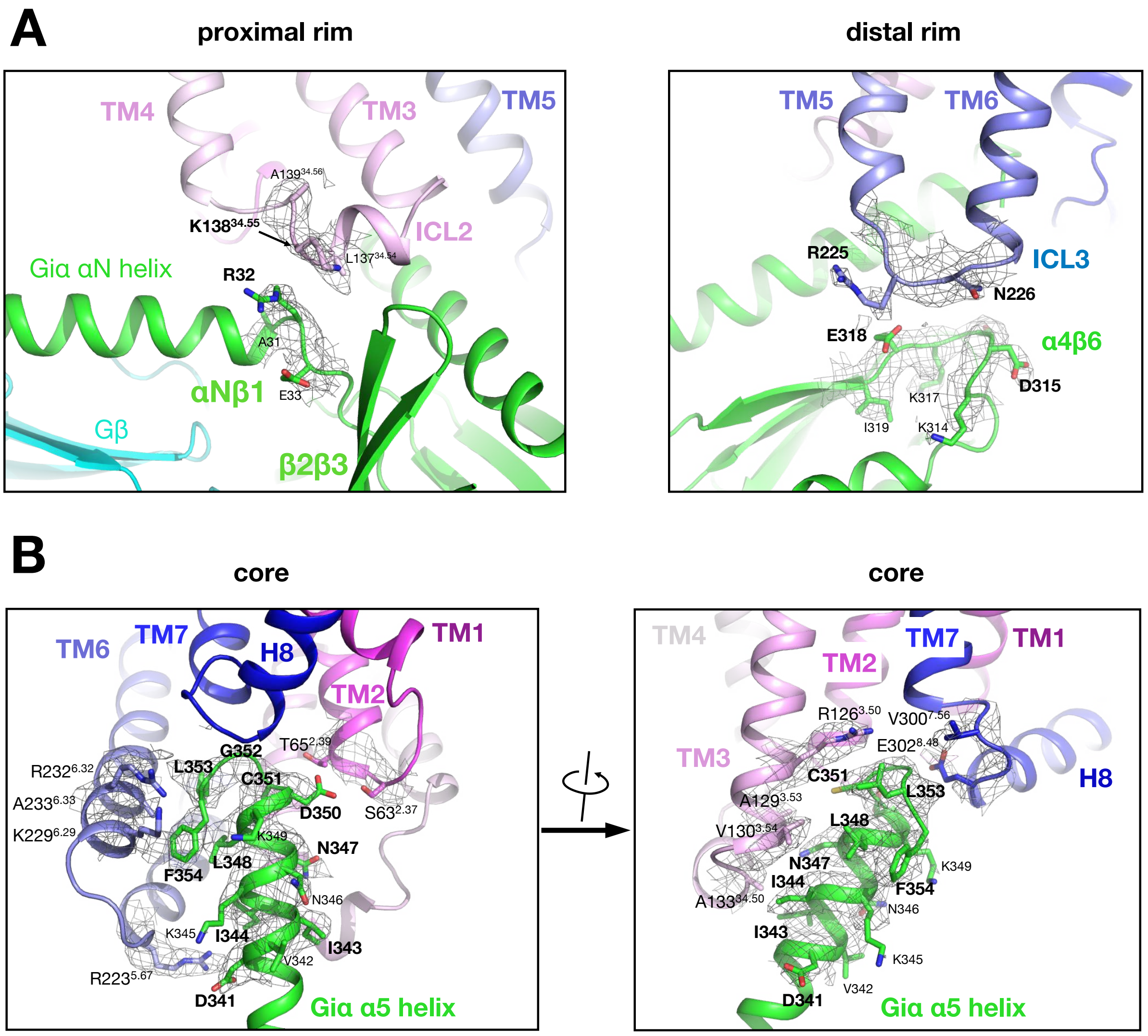


**Figure S11. Electron density at the contact interface between CCR5 and Giα.** (A) Left: contacts at the proximal rim of the interface, formed by ICL2 of CCR5 and the αNβ1/β2β3 loops of Giα. The mesh depicts the cryo-EM map contoured at a 7.5 σ cut-off for residues 137-139 of CCR5 and 31-33 of Giα. Right: contacts at the distal rim of the interface, formed by ICL3 of CCR5 and the α4β6 loop of Giα. The mesh depicts the cryo-EM map contoured at a 7.5 σ cut-off for residues 224-227 of CCR5 and 314-319 of Giα. (B) Contacts at the core of the interface, formed by ICL1 and ICL4 the cytoplasmic ends of TM3, TM5, and TM6 of CCR5 and the α5 helix of Giα. CCR5 residues within 4 Å of the Giα α5 helix and residues 341-354 of the Giα α5 helix are displayed as sticks with the grey mesh depicting the cryo-EM map at a 7.5 σ cut-off. Residues on the α5 helix residing within 4 Å to CCR5 are labelled in bold.

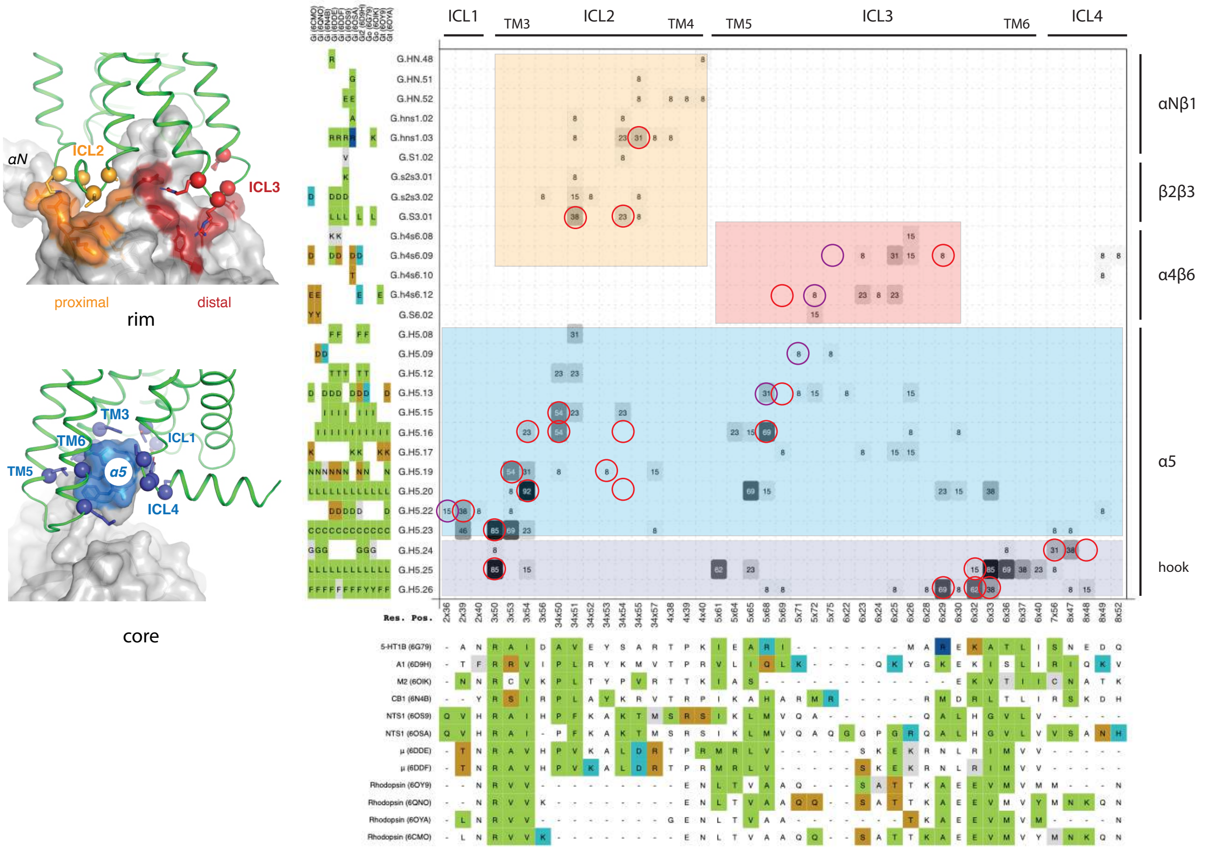


**Figure S12. Receptor/Gα protein contact map of currently available GPCRs/G$_i$ complexes.** The map was obtained from www.gpcr.org. GPCR residues (in ICL1, ICL2, ICL3, and ICL4 and in the cytoplasmic ends of TM3, TM4, TM5, and TM6) are in the X-axis and Gα$_i$ residues (in the αNβ1, β2β3, α4β6 loops and in the α5 helix) are in the Y axis. The number of contacts at each position (in all available complexes) is depicted in the map in a grey scale (light grey: few contacts at this position across complexes; dark grey: many contacts). Red circles represent the contacts present in the CCR5/[6P4]CCL5 complex; purple circles represent contacts present in our complex but shifted by one residue compared to other complexes. Next to the axes, the sequence composition in each GPCR or Gα$_i$ position is shown, with residues colored by their physico-chemical nature. The proximal rim of the interface is highlighted in orange, the distal rim in red, and the core in light (base of α5) and dark (hook of α5) blue. These interfaces are shown on the 3D molecular structure at the left. At the proximal rim of the interface (orange), the CCR5/[6P4]CCL5 features several of the 'expected' contacts (red circles). On the other hand, the distal rim of the interface presents contacts absent in other complexes (purple circles). At the core of the interface, many of the expected contacts are present (red circles), while there are several contacts unique for the CCR5/[6P4]CCL5 complex. While there is certain variability in the precise composition of the interfaces, the CCR5/[6P4]CCL5 complex is overall similar to existing complexes, with some variability at the distal rim and the α5 hook.

**A**

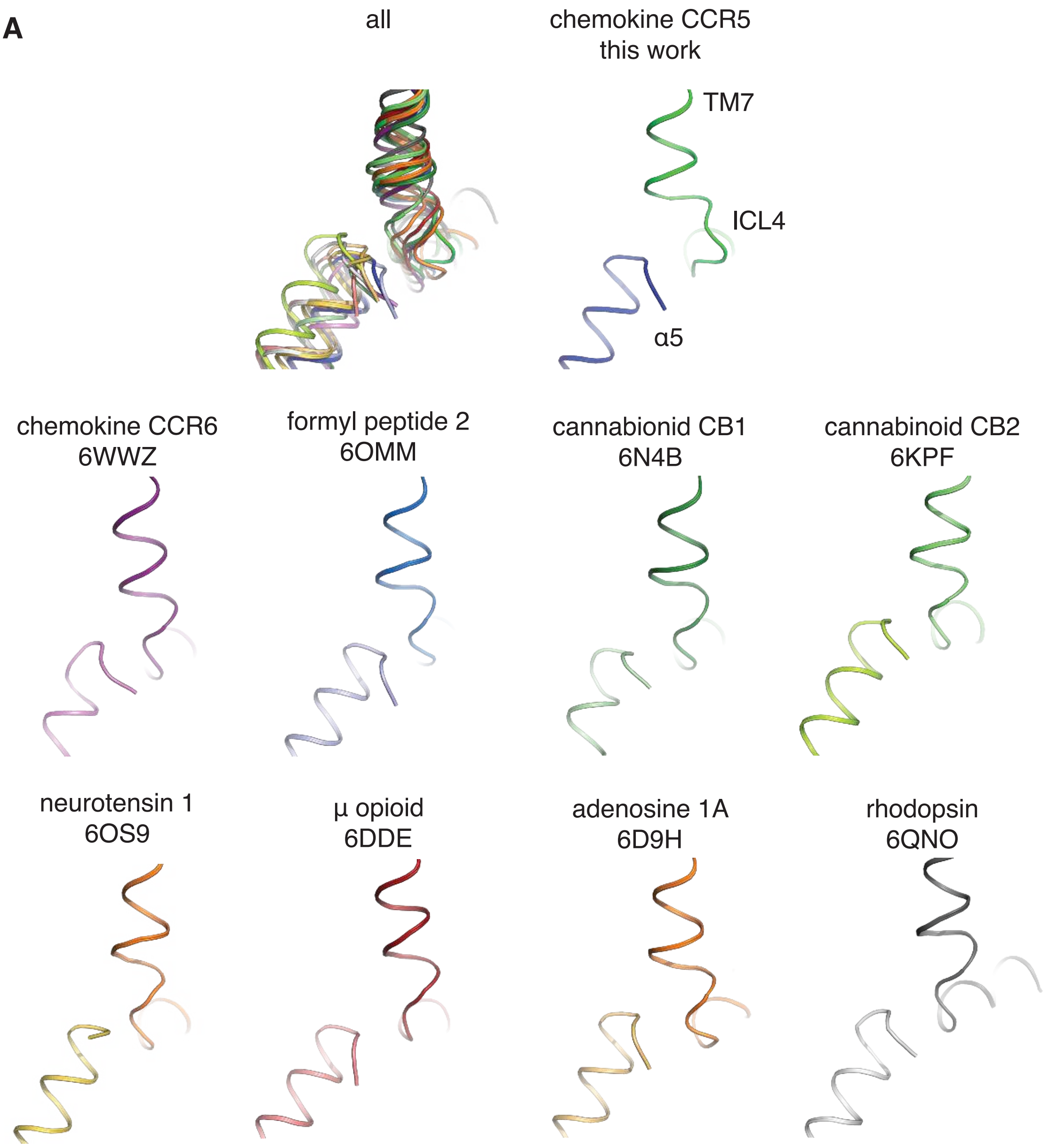


**B**

```
>ccr5       AMQVTETLGMTHCCINPIIYAFVGEKFRN
>ccr6       TKTVTEVLAFLHCCLNPVLYAFIGQKFRN
>fp2-6omm   LVNPTSSLAFFNSCLNPMLYVFVGQDFRE
>cb1-6n4b   VFAFCSMLCLLNSTVNPIIYALRSKDLRH
>cb2-6kpf   AFAFCSMLCLINSMVNPVIYALRSGEIRS
>nt1-6os9   FYMVTNALFYVSSTINPILYNLVSANFRH
>mop-6dde   SWHFCIALGYTNSCLNPVLYAFLDENFKR
>aa1r-6d9h  LTYIAIFLTHGNSAMNPIVYAFRIQKFRV
>rho-6qno   FMTIPAFFAKTSAVYNPVIYIMMNKQFRN
            TM7                 ICL4  HX8
```

**Figure S13. Structure and sequence of ICL4 in available GPCRs in complex with $G_{i/o}$.** (A) A structural alignment (on the GPCR transmembrane bundle) of GPCR/$G_{i/o}$ structures reveals the plasticity of ICL4. (B) A sequence alignment of ICL4 in GPCRs bound to $G_{i/o}$ shows the sequence variability in this region.

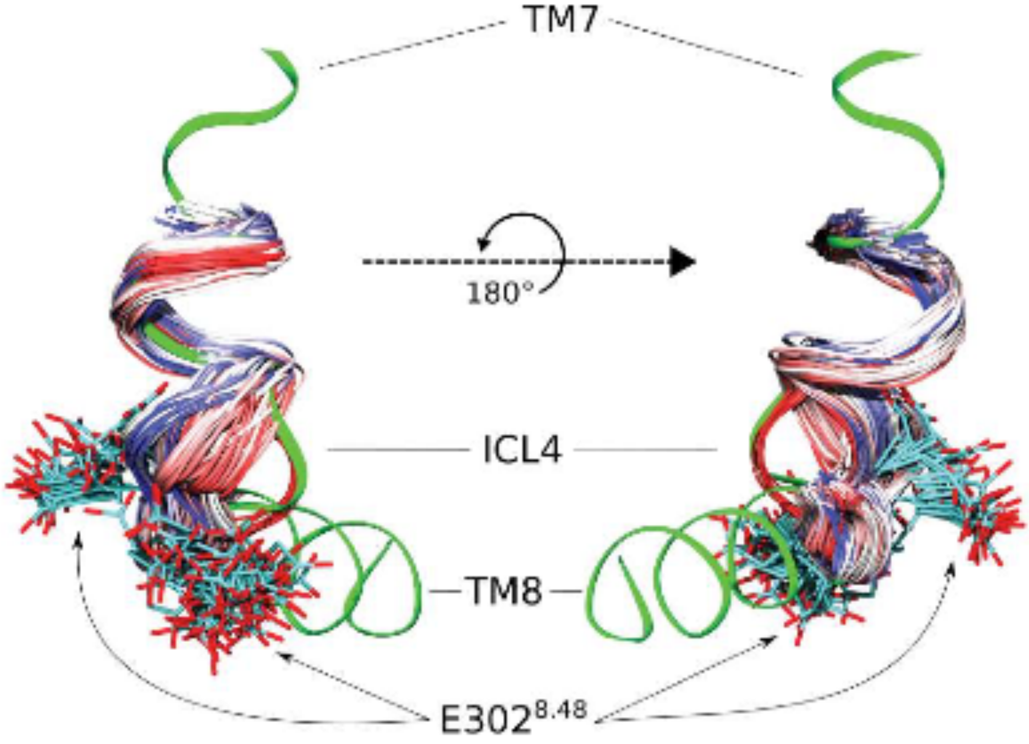


**Figure S14. Conformational dynamics of ICL4 in MD simulations of the [6P4]CCL5•CCR5 complex.** The CCR5 TM7-ICL4-HX8 junction in the cryo-EM structure described in this work is shown as a green cartoon. A representative conformational ensemble of CCR5 residues 296 to 304 from our molecular dynamics simulations (in the absence of bound $G_{\alpha i}$) is depicted as colored cartoons. Time evolution during the simulation is represented in a blue-to-red color gradient (blue – beginning of the simulation; red – end of the simulation). The side chain of residue $E302^{8.48}$ is shown as sticks. Our simulations indicate that the local structure of ICL4 is plastic, adapting to changes in the environment. In the absence of the bound $G_{\alpha i}$, ICL4 and the side chain of $E302^{8.48}$ adopt two preferred conformations.

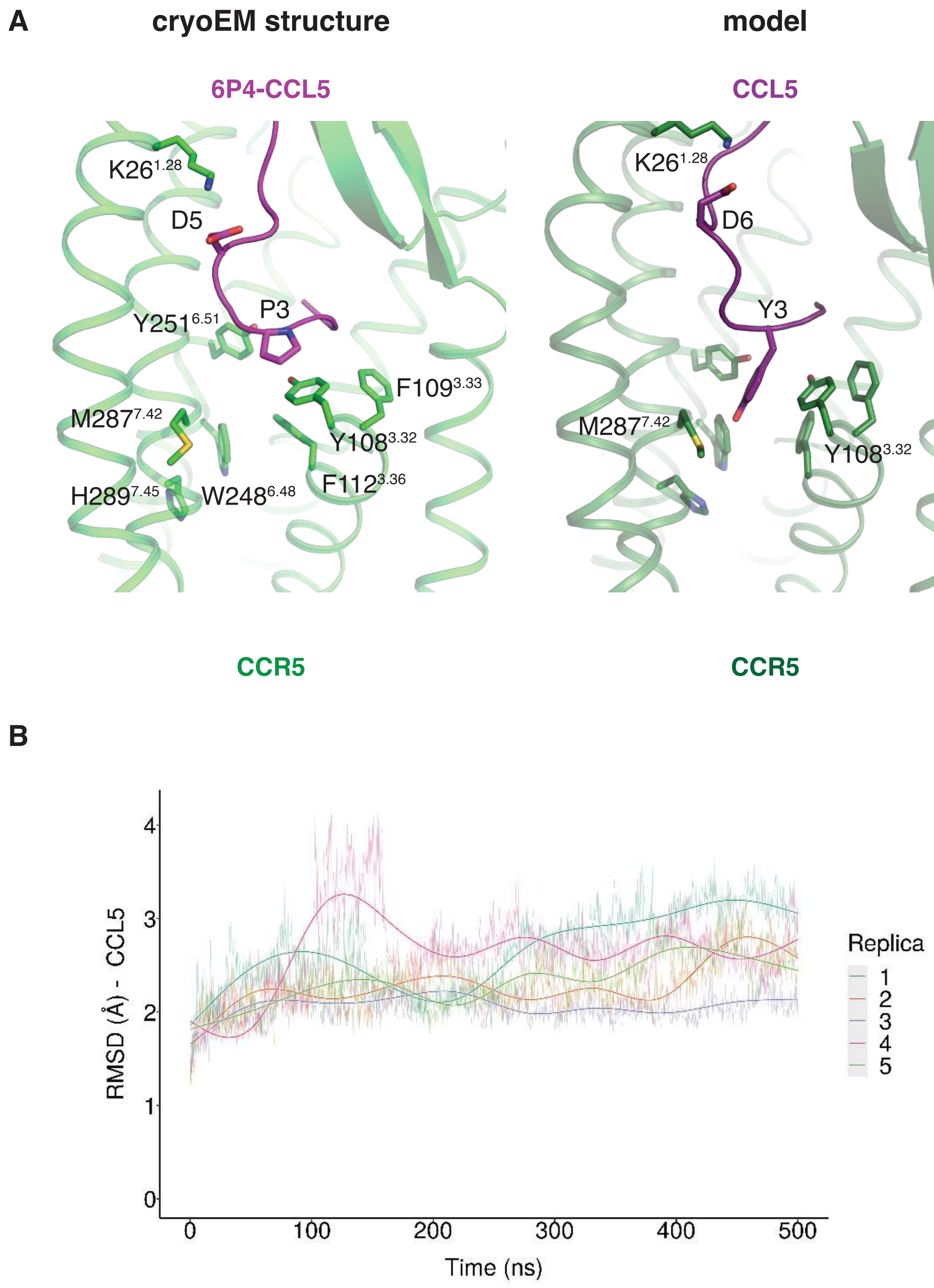


**Figure S15. Model of wild type CCL5 bound to CCR5.** (A) Left: cryo-EM structure of the [6P4]CCL5•CCR5 complex. D5 in the chemokine interacts with K26$^{1.28}$ in the receptor stabilizing the conformation of the CCL5 N-terminus. P3 in the chemokine pushes on M287$^{7.42}$ and Y108$^{3.32}$ in the aromatic connector. Right: model of wild type CCL5 in complex with CCR5. In this case, residues D6 and Y3 in the chemokine may play the same role as D5 and P3 in our structure. (B) Root mean square deviation (RMSD) of the modeled CCL5 in MD simulations of the CCR5•CCL5 complex. Each line represents an independent simulation replica, and the overlaid bold lines represent smoothed averages. The data show that the modeled pose of CCL5 is stable during the simulations.

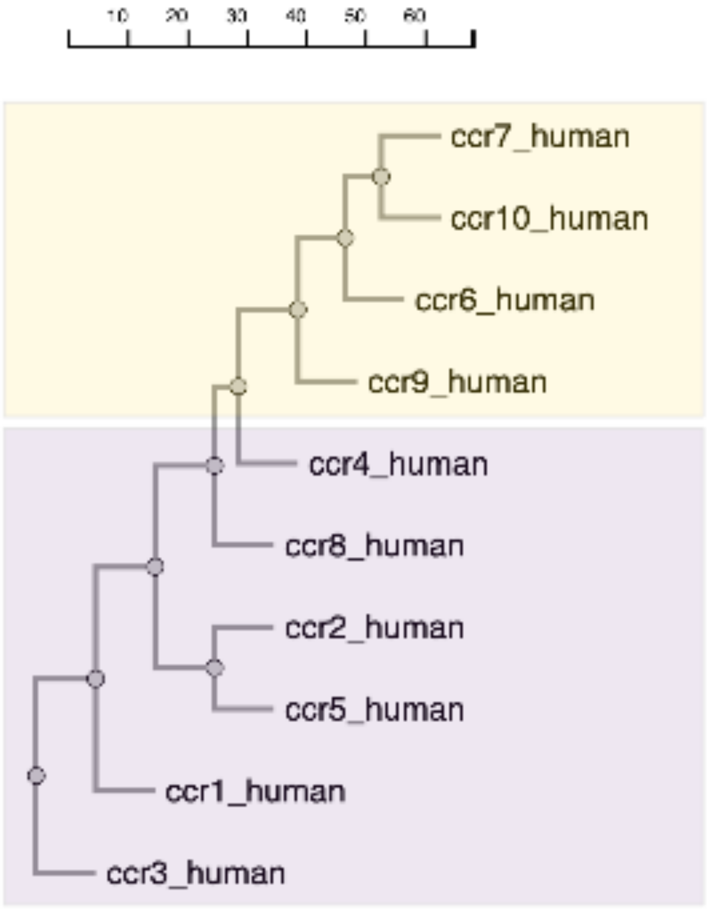


**Figure S16. Phylogenetic tree of CC chemokine receptors.** The tree was obtained at www.gpcrdb.org (see Methods), and shows how CC chemokine receptors containing $W^{6.48}$ (CCR1-5 and CCR8; magenta) and those containing $Q^{6.48}$ (CCR6, CCR7, CCR9, and CCR10; yellow) form distinct subgroups (see Figure 5 in the main text).
