## Supplementary Video Legends for "Structural basis of the activation of the CC chemokine receptor 5 by a chemokine agonist"

**Supplementary Video S1. 3D variability analysis (3DVA) of the [6P4]CCL5•CCR5•Gi•Fab16 cryo-EM map.** The analysis resulted in variability components 0, 1 and 2 displayed in the videos S1a, S1b and S1c. The following motions can be perceived in the 3DVA: bending of the C-terminal helix of [6P4]CCL5 towards CCR5; variation of the insertion depth of the [6P4]CCL5 N-terminus in the CCR5 binding pocket; opening of the binding pocket of CCR5 by the rearrangement of the 7TM bundle; correlated motions of $G_i$ and Fab16; twisting and contraction of the transmembrane helix bundle of CCR5; and elongation of TM3, which maintains the contact of ICL2 with $G\alpha_i$. The dynamic nature of GPCRs produces heterogeneous density maps in Cryo-EM studies, which leads to locally blurred density maps. 3DVA allows to visualize this dynamic behavior of the complex and reveals several modes of possible functional significance which cannot be described by one density map and single structure.

**Supplementary Video S2. Simulation of the interaction between the sulfated CCR5 N-terminus and [6P4]CCL5.** CCR5 is displayed as white cartoons with its N terminus in green. Sulfated tyrosines Y10 and Y14 are shown as sticks (cyan carbon atoms) and [6P4]CCL5 is shown as a pink surface.

**Supplementary Video S3. Representative simulation of the CCL5•CCR5 model.** CCR5 is represented as green cartoons, and CCL5 in magenta. Key residues are shown as cyan sticks. In the molecular dynamics trajectory, residue Y3 of CCL5 (at the center of the image) engages the aromatic connector of CCR5 ($Y108^{3.32}$, $F109^{3.33}$, $F112^{3.36}$ (center left), and $Y251_{6.51}$ (center right)). Residue E6 of CCL5 engages K26 of CCR5 forming a salt bridge (top). Both features are preserved during the MD simulations.
